# STARsolo: accurate, fast and versatile mapping/quantification of single-cell and single-nucleus RNA-seq data

**DOI:** 10.1101/2021.05.05.442755

**Authors:** Benjamin Kaminow, Dinar Yunusov, Alexander Dobin

## Abstract

We present *STARsolo*, a comprehensive turnkey solution for quantifying gene expression in single-cell/nucleus RNA-seq data, built into RNA-seq aligner *STAR*. Using simulated data that closely resembles realistic scRNA-seq, we demonstrate that *STARsolo* is highly accurate and significantly outperforms pseudoalignment-to-transcriptome tools. *STARsolo* can replicate the results of, but is considerably faster than *CellRanger*, currently the most widely used tool for pre-processing scRNA-seq data. In addition to uniquely mapped reads, *STARsolo* takes account of multi-gene reads, necessary to detect certain classes of biologically important genes. It has a flexible cell barcode processing scheme, compatible with many established scRNA-seq protocols, and extendable to emerging technologies. *STARsolo* can quantify transcriptomic features beyond gene expression, which we illustrate by analyzing cell-type-specific alternative splicing in the Tabula Muris project.

## 1 Introduction

Single-cell RNA-sequencing (scRNA-seq) has revolutionized the analysis of complex biological systems, providing unprecedented insight into transcriptomic profiles of individual cells and enabling high-throughput characterization of cellular composition of tissues, tumor microenvironments, developmental programs and microbial communities. A multitude of bioinformatic tools have been developed to perform a variety of scRNA-seq analyses, such as identifying known and novel cell types, pin-pointing cell type markers, reconstructing gene regulatory networks, and inferring developmental trajectories [1].

Single-cell RNA-seq technologies generate read sequences containing three key pieces of information: cDNA fragment that identifies the RNA transcript; cell barcode (CB) that identifies the cell where the RNA was expressed; and Unique Molecular Identifier (UMI) that identifies the specific RNA molecule [2]. The foundational stage of the scRNA-seq analyses is mapping/quantification, which involves four main steps: (i) mapping the cDNA fragments to a reference, (ii) assigning reads to genes, (iii) assigning reads to cells (cell barcode demultiplexing), and (iv) counting the number of unique RNA molecules (UMI deduplication). The outcome of this procedure is the gene/cell expression matrix, which contains the counts of RNA molecules in each cell for each gene. The mapping/quantification accuracy is of the utmost importance, since it serves as the starting point for all the downstream analyses. This procedure is also computationally demanding, as it requires processing hundreds of millions of reads generated in a typical scRNA-seq experiment.

A number of mapping/quantification tools have been developed recently, and they can be classified in two broad categories. The fully modular tools [3–6] use standard RNA-seq aligners that are not aware of the intrinsics of the scRNA-seq protocol, while protocol-specific CB/UMI processing happens in a separate step. The advantage of this approach is its high flexibility in selecting the aligner, read-to-gene assignment and CB/UMI processing methods. However, since the communication between different steps happens via generic file formats (e.g. FASTQ and BAM), this approach typically results in low processing speed and overtaxing of computing infrastructure. The next wave of tools, *Alevin* [7] and *Kallisto-bustools* [8], significantly increased the overall computational efficiency owing to a tight integration between alignment and CB/UMI processing steps. To achieve high performance without sacrificing modularity, these tools utilize efficient intermediate file formats [9, 10].

In this manuscript we present *STARsolo* [11], a comprehensive turnkey solution for mapping/quantification of scRNA-seq data. *STARsolo* is built directly into the RNA-seq aligner STAR [12], which is widely used for mapping bulk RNA-seq data. In *STARsolo*, read mapping, read-to-gene assignment, cell barcode demultiplexing and UMI collapsing are tightly integrated (Methods 5.1), avoiding input/output bottlenecks and boosting the processing speed. Importantly, *STARsolo* performs read alignment to the full genome, resulting in a higher accuracy compared to the alignment to transcriptome only.

To evaluate the mapping/quantification accuracy we developed a strategy for simulating scRNA-seq reads based on real RNA-seq data (Methods 5.2). We used simulated data to compare *STARsolo* with *Kallisto* and *Alevin*, and demonstrated high accuracy of *STARsolo* gene quantification (Results 2.2, 2.4). On the other hand, we found a significant reduction of gene quantification accuracy for tools that utilize pseudoalignment-to-transcriptome, resulting in false positive detection of thousands of genes (Results 2.3).

To compare computational efficiencies of different mapping/quantification algorithms, we benchmarked the processing speed and Random Access Memory (RAM) usage of *STARsolo*, *CellRanger*, *Alevin* and *Kallisto* on a real dataset (Results 2.7). While pseudoalignment-to-transcriptome tools are the fastest and require least RAM, *STARsolo* is the fastest among the highly accurate methods. Moreover, STAR uses less RAM and is much faster than pseudoalignment-to-transcriptome for more complicated analyses such as RNA Velocity [13] calculations or single-nucleus RNA-seq quantification.

*STARsolo* is designed to be a drop-in replacement for the *CellRanger* [14], a proprietary tool from 10X Genomics company, which presently is the dominant commercial scRNA-seq platform. *CellRanger* uses *STAR* for mapping reads to the reference genome, while the other steps of the gene quantification pipeline are performed with its own proprietary algorithms. With an appropriate choice of parameters, *STAR-solo* can generate a gene/cell count matrix almost identical to *CellRanger*’s (Results 2.5). The other tools deviate significantly from *CellRanger*, both in overall gene expression quantification and, crucially, in differential expression of cell-type-specific marker genes (Results 2.6).

Unlike *CellRanger*, which is hard-coded for analyzing 10X Genomics products, *STARsolo* features a flexible CB/UMI processing scheme, which is compatible with many established protocols and can be easily extended to emerging technologies. Another important advantage of *STARsolo* is the ability to quantify reads that map to multiple genes (Results 2.4). The importance of including multi-gene reads had been discussed before for bulk [15–19] and single-cell RNA-seq data [7, 10, 20]. Furthermore, *STARsolo* is capable of investigating transcriptomic features beyond standard gene expression, which we illustrate by analyzing cell-type-specific alternative splicing in the Tabula Muris scRNA-seq dataset (Results 2.8).

## 2 Results

We compared *STARsolo* performance on simulated and real data with several existing tools: *CellRanger* [14], *Alevin/Alevin-fry* [7, 10, 21] and *Kallisto/Bustools* [8, 9, 22]. *CellRanger* is a de facto standard for analyzing 10X Genomics scRNA-seq data, while *Kallisto* and *Alevin* use light-weight alignment-to-transcriptome algorithms which are profoundly different from STAR’s aligment to the full genome. For *Alevin*, we used 4 different alignment modes: pseudoalignment and selective alignment to transcriptome only, as well selective alignment with partial and full genome decoys. As we will see below, these options have a strong impact on the mapping/quantification accuracy.

### 2.1 Simulating scRNA-seq reads

To quantitatively gauge the accuracy of mapping/quantification tools, we need to know the true gene expression values, which can only be accomplished by simulating scRNA-seq data. We note that most published scRNA-seq simulation methods (e.g. [23–26]) simulate the gene/cell count matrix but not the scRNA-seq read sequences that are required to test the mapping/quantification approaches.

Typically, bulk RNA-seq reads are simulated using a predefined distribution of reads across the RNA transcripts, modeled after a distribution observed in real RNA-seq data. Such an approach, however, is unfeasible for scRNA-seq reads, owing to a non-trivial distribution of reads through the transcript. To bypass this complication, we propose a simulation strategy that closely reflects real scRNA-seq reads (see Methods 5.2 for details). For a selected real scRNA-seq dataset, we map the cDNA reads to the transcriptome and full genome simultaneously, using the highly accurate *BWA-MEM* [27] aligner. For each read the “true locus” is selected randomly from the top-scoring *BWA-MEM* alignments, with the preference given to the transcript over genomic alignments. Then, the cDNA read sequence is simulated from the true locus sequence, with substitution errors added at a constant mismatch error rate of 0.5%, typical for good quality Illumina sequencing. Each read is given its real CB and UMI, if the CB is present in the CB-passlist. UMIs for the reads whose “true locus” is a transcript are counted towards the corresponding genes to generate the true gene/cell count matrix.

Our simulation approach does not deal with the full complexity of real scRNA-seq data: for instance, it avoids the issues of CB and UMI error correction. However, by simulating reads from a realistic distribution of transcriptomic and genomic loci, we can evaluate the accuracy of the most crucial steps of the algorithms: read mapping and read-to-gene assignment.

### 2.2 Simulations without multi-gene reads

One of the important, not yet fully solved questions of both bulk [15–19] and scRNA-seq [7, 10, 20] analyses is how to deal with multi-mapping reads. The definition of a multi-mapping read is somewhat inconsistent throughout the literature. In studies where reads are mapped to the whole genome, multimappers are defined as reads that map to two or more genomic loci. Yet if reads are mapped to the transcriptome, the multimappers are defined as reads that map to two or more transcripts. For scRNA-seq analyses, we count the number of reads per gene, hence what matters is whether a read maps to one or multiple genes. For clarity, we will call reads that map to two or more genes “multi-gene reads”. Note that a multi-gene read can map to only one genomic locus where two or more genes overlap, or it can map to multiple genomic loci that contain multiple genes. On the other hand, a read that maps to multiple genomic loci, but overlaps a gene in only one locus, is considered a “unique-gene” read.

To investigate the effect of multi-gene reads on the mapping quantification accuracy, we first performed simulations *excluding multi-gene reads*. Accordingly, we ran all tools with options that discard multi-gene reads (see Supp. Table 1 for tools’ parameters).

We derived the simulated dataset from the 10X Peripheral blood mononuclear cell dataset generated by the 10X Chromium v3 protocol (10X-pbmc-5k, Supp. Table 1). This dataset is a representative of typical 10X libraries, containing 384 million reads for ~5000 cells, with ~8000 UMIs per cell on average. This dataset will be used throughout this manuscript for both simulated and real data comparisons.

All metrics here (and everywhere in this study) are calculated for a set of “filtered” cells, common for all tools. For simulations, this set is created by intersecting the top 5000 cells by their true UMI counts with raw cells from *STARsolo*, raw cells from *Kallisto* (which does not perform cell filtering), and filtered cells from *Alevin* (which does not output unfiltered cells).

Figure 1A shows the overall Spearman correlation coefficient between true simulated gene/cell counts and those produced by each tool (see Supp. Fig. S1A for the full correlation matrix). The correlation is calculated between the elements of the gene/cell count matrices which are expressed (i.e. count ≥1 UMIs) in *either* the simulation or tool quantification. We can see that *STARsolo* achieves a nearly perfect agreement with true counts *R*≈0.99, while the pseudoalignment-to-transcriptome algorithms (*Kallisto* and *Alevin_sketch*) show the lowest correlation (*R*≈0.77 and 0.8 respectively). *Alevin*’s accuracy increases as its alignment algorithm changes from pseudoalignment to selective alignment, and it improves further when a partial and, finally, full genome decoy is used in addtion to the transcriptome. Ultimately, *Alevin_full-decoy* attains correlation with truth as high as *STARsolo*.

**Figure 1:**
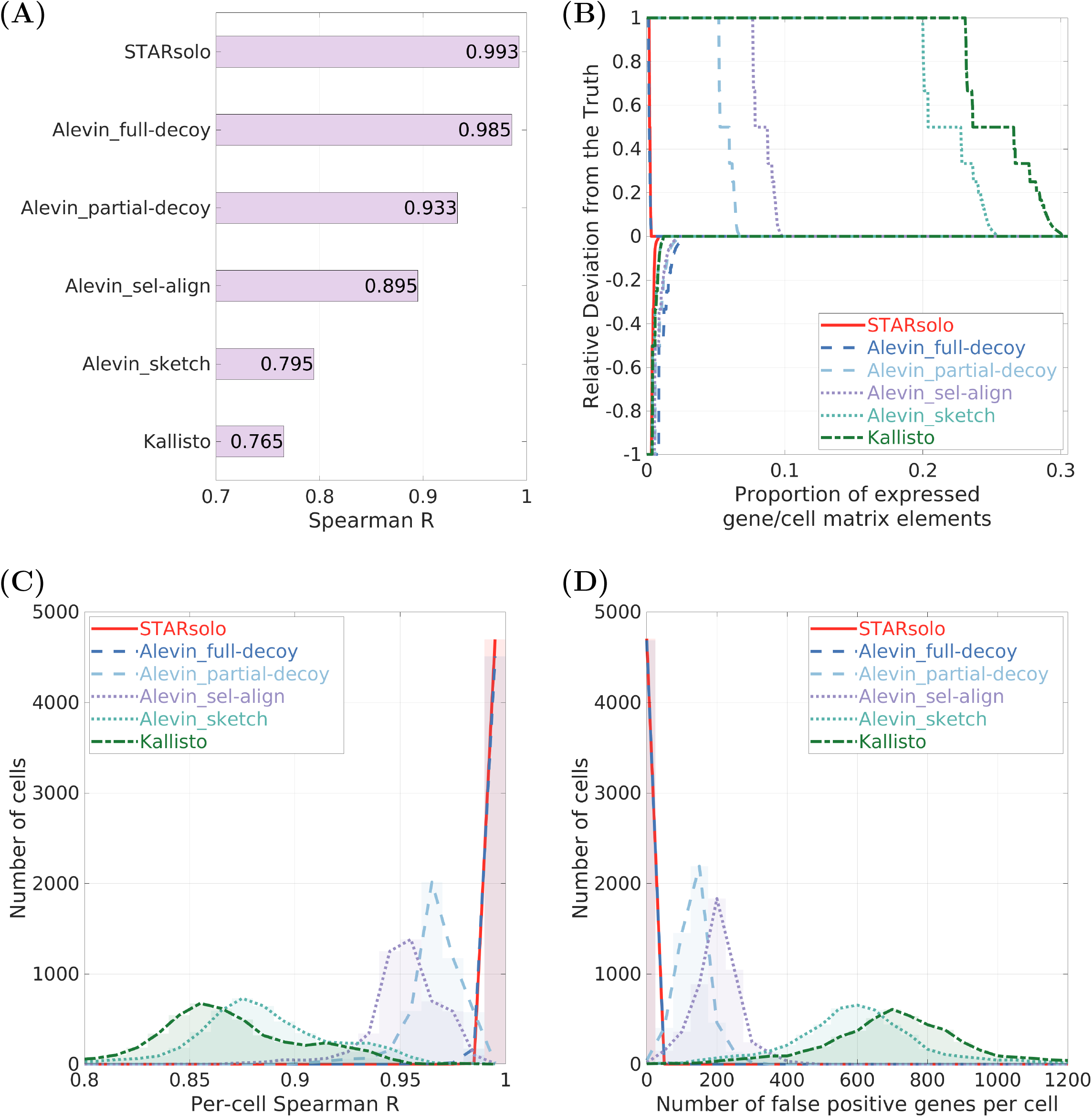
Simulations **without multi-gene reads** (Results 2.2). **(A):** Spearman correlation coefficient R vs simulated truth for gene/cell matrix elements. **(B):** Relative Deviation for gene/cell matrix elements with respect to the true counts. **(C):** Histogram of per-cell Spearman correlation coefficients vs truth. **(D):** Histogram of the number of false positive genes per cell: genes that were *not* expressed in the simulation, but were detected by each tool.

Another metric, Relative Deviation (*RD*) of a tool’s gene/cell counts with respect to the simulated truth, is defined as *RD* = (*C*_*tool*_ – *C*_*true*_)/*max*(*C*_*tool*_, *C*_*true*_) and is presented in Figure 1B (Supp.Fig. S1B shows Mean Absolute RD). The negative values of RD correspond to false negatives, i.e. a tool not detecting gene/cell counts that were simulated. The positive values of RD correspond to false positives, i.e. gene/cell counts that were not present in the simulated data but were predicted by a tool. Figure 1B shows that all tools have low false negative rates (i.e. high sensitivity), with only several percent of genes/cells having *RD*<0. Strikingly, while *STARsolo* and *Alevin_full-decoy* show a very low false positive rate, *Kallisto* and *Alevin_sketch* predict false expression for >20% of gene/cell matrix elements. As before, the *Alevin* performance improves as it moves away from pseudoalignment-to-transcriptome to the selective alignment with full genome decoy.

The third metric is the per-cell Spearman correlation coefficient between each tool and the truth, calculated in each cell (Figure 1C, cell-averaged 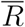 shown in Supp.Fig. S1B). In this calculation we include only those genes that are expressed in any of the cells in either the simulation or tool matrices. *STARsolo* and *Alevin_full-decoy* show high congruity to the ground truth for almost all cells 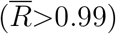, while other tools yield a wide distribution of lower correlations, with *Kallisto* and *Alevin_sketch* again showing the worst performance (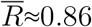 and 0.88 respectively).

### 2.3 Pseudoalignment-to-transcriptome predicts expression for thousands of non-expressed genes

To further investigate the impact of algorithmic choices on mapping/quantification accuracy, we compared the genes predicted to be expressed in each cell by each tool to the simulated truth. Figure 1D shows the distribution of the number of false positive genes per cell, i.e. genes that were not expressed in the simulated data, but were predicted by a tool. While *STARsolo* and *Alevin_full-decoy* call only a few false positive genes per cell (~5 on average), the pseudoalignment-to-transcriptome tools predict expression for a vast number of non-expressed genes, on average ~600 for *Kallisto* and ~700 for *Alevin_sketch* in each cell, which constitutes a substantial proportion of all genes expressed in a cell (≈21% and 24% respectively, Supp.Fig. S1B).

We believe that this severe overprediction of gene expression by the pseudoalignment-to-transcriptome tools is caused by the reads that originate in the non-exonic regions of the genome (intronic and intergenic), but are forced to map to the sequence-similar regions of the transcriptome. Intronic reads are very abundant in single-cell RNA-seq data [13], likely owing to non-specific priming of poly-dT reverse transcription primers. The full and partial decoy options for *Salmon*/*Alevin* were developed to specifically mitigate the adverse effect of non-transcriptomic reads [28], which explains their significant performance improvement over pseudoalignment-to-transcriptome-only approach.

To confirm this hypothesis we excluded the non-exonic reads from our simulated data. This indeed resulted in a significant accuracy improvement for the pseudoalignment-to-transcriptome tools (see Figure S2). In reality, however, current droplet-based single cell RNA-seq technologies contain a large (~20-40%) proportion of intronic reads [13], making pseudoalignment-to-transcriptome-only approach unreliably inaccurate for such technologies.

### 2.4 Simulation with multi-gene reads

Since multi-gene reads constitute a relatively small proportion of all reads, they are discarded in many bulk and single-cell RNA-seq pipelines. However, their exclusion may conceal expression of important genes and gene families [15–19]. To include the multi-gene reads, the expectation-maximization algorithm was introduced in *Alevin*, and its impact on gene detection and quantification was investigated [7, 20]. A similar option was also implemented in *Kallisto* [8].

We implemented several options in STARsolo for multi-gene reads recovery, ranging from a simple uniform distribution of multi-gene reads among their genes, to a full expectation-maximization model, which was used for the comparisons below (Methods 5.1.7). In Figure 2, we present the accuracy metrics for simulations that include multi-gene reads. All tools were run with their multi-gene options (Supp. Table 1). Only gene/cell count matrix elements ≥0.5 were considered expressed.

**Figure 2:**
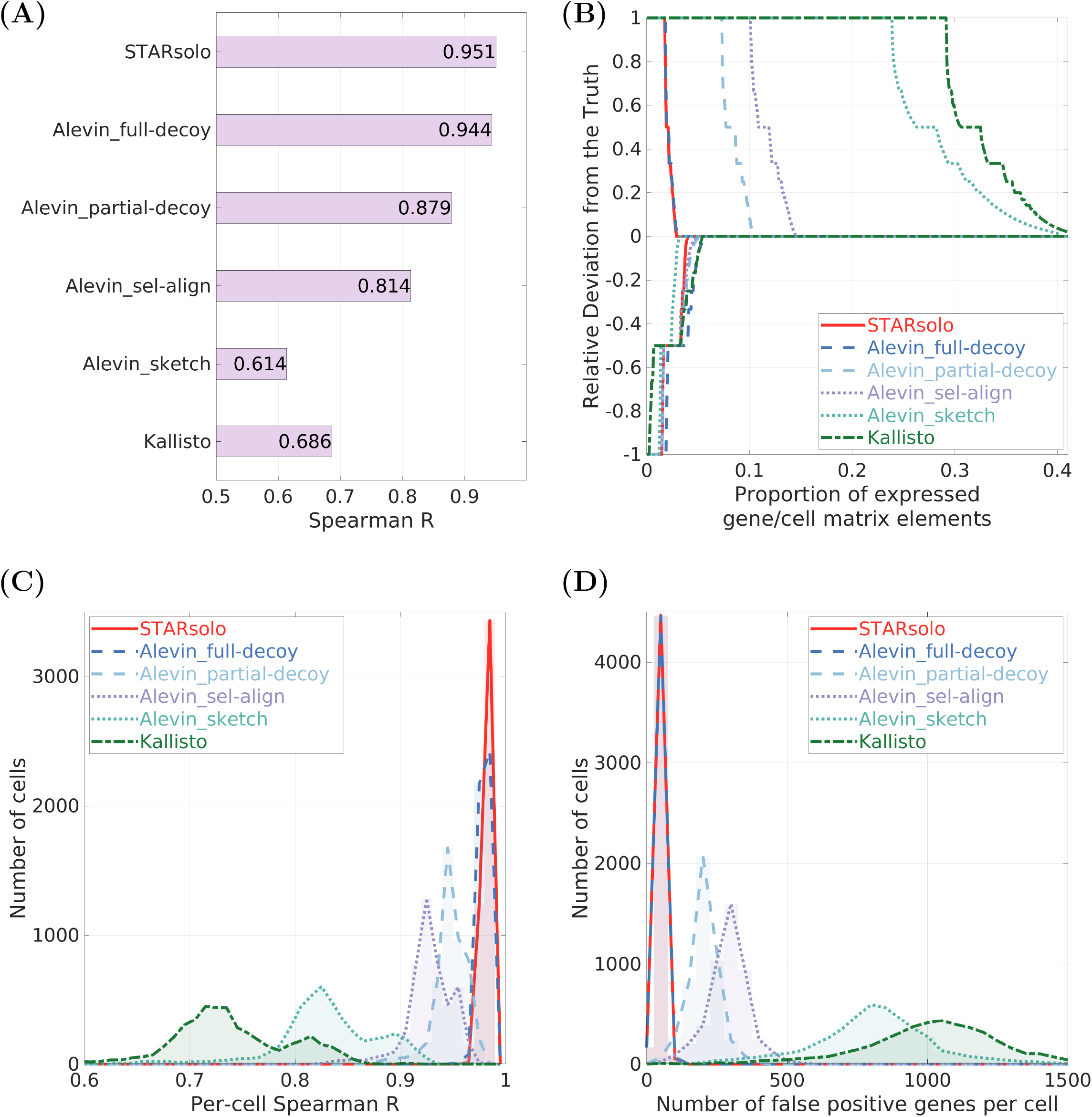
Simulations **with multi-gene reads** (Results 2.4). **(A):** Spearman correlation coefficient R vs simulated truth for gene/cell matrix elements. **(B):** Relative Deviation for gene/cell matrix elements with respect to the true counts. **(C):** Histogram of per-cell Spearman correlation coefficients vs truth. **(D):** Histogram of the number of false positive genes per cell: genes that were *not* expressed in the simulation, but were detected by each tool.

Comparing Figures 1A and 2A, we see that when simulations include multi-gene reads, the genes/cells Spearman correlation to the true counts is reduced for all tools. For the most accurate tools, *STARsolo* and *Alevin_full-decoy*, it drops from *R*≈0.99 to 0.95. For the pseudoalignment-to-transcriptome tools the reduction of correlation is larger: from *R*≈0.86 to 0.69 for *Kallisto*, and from *R*≈0.9 to 0.61 for *Alevin_sketch*.

Multi-gene reads also reduce per-cell Spearman correlation with the true counts (Figures 2D vs 1D) and diminish the precision of gene detection (Figures 2C vs 1C). While *STARsolo* and *Alevin_full-decoy* keep false positive genes in check, the pseudoalignment-to-transcriptome tools further inflate the number of false positive genes, on average from ~700 to ~1,100 for *Kallisto*, and from ~600 to ~800 for *Alevin_full-decoy*.

The reason for this reduction in accuracy becomes clearer if we compare the relative deviation (RD) from the true counts for simulations with and without multi-gene reads (Figures 2B vs 1B). We see that the proportion of false negatives (negative RD) increases when multi-gene reads are included, which indicates that multi-gene algorithms cannot recover correctly all multi-gene reads. At the same time, the proportion of false positives (positive RD) also increases, indicating that the algorithms are assigning some of the multi-gene reads to the non-expressed genes.

To further elucidate the impact of multi-gene recovery, we run the tools with and without multi-gene recovery options (Figure 3). Without multi-gene recovery, *STARsolo_mult:No* and *Alevin_full-decoy_mult:No* have a very low false positive rate, but, owing to exclusion of multi-gene reads, yield a higher (~5%) false negative rate. Turning on multi-gene read recovery reduces the false negative rate, at the cost of some increase the false positive rate, resulting in an overall accuracy improvement. While this improvement in accuracy is relatively small, it can have serious implications for biological inferences derived in downstream analyses. For instance, it may reveal expression of functionally important gene paralogs specific to certain cell types.

**Figure 3:**
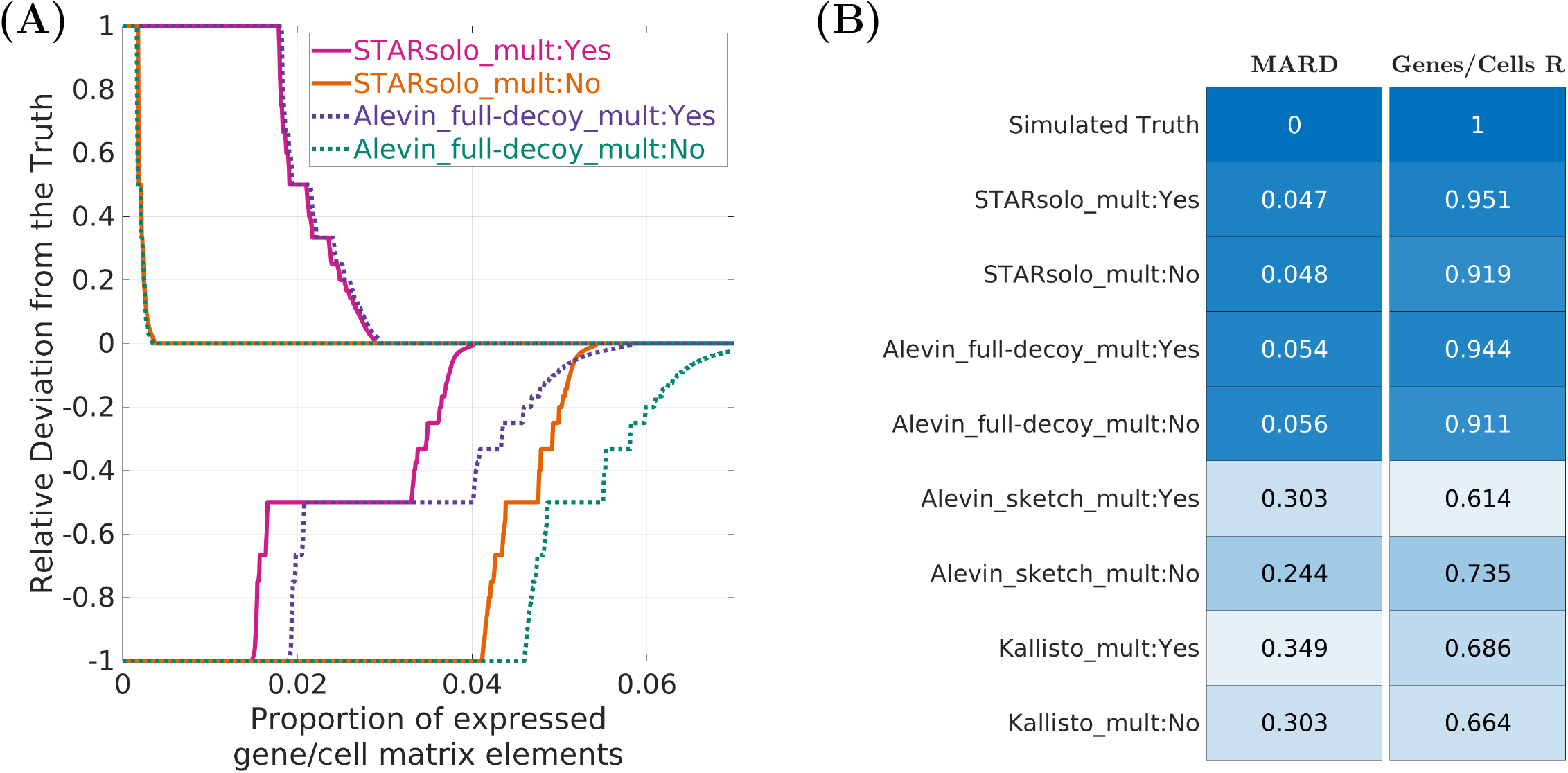
Simulations **with multi-gene reads**: multi-gene vs no-multi-gene algorithms (Results 2.4). **(A):** Relative Deviation for gene/cell counts with respect to true counts. **(B):** Mean Absolute Relative Deviation and Spearman correlation for gene/cell matrix elements with respect to the true counts.

### 2.5 Mapping/quantification comparison on real data

To evaluate tools’ performances on real scRNA-seq data, we used the 10X Peripheral blood mononuclear cell dataset generated by the 10X Chromium v3 protocol (10X-pbmc-5k, the same dataset as was used to generate the simulated data). To compare all tools on the same set of cells, we computed the intersection between *CellRanger* filtered cells (5026), *STARsolo* and *Kallisto* raw (unfiltered) cells, and *Alevin* filtered cells. The resulting intersection contains 4655 cells. In real user experience, the cell calling (filtering) will produce different sets of cells for *Alevin* and *Kallisto*, which will further amplify their differences to *CellRanger*. *STARsolo*, on the other hand, can perform cell filtering identically to *CellRanger* (Methods 5.1.8).

Figures 4 and S4 show the same accuracy metrics as were used for the simulated data, but each tool is compared to the *CellRanger* instead of the simulated truth, since the latter is not available for the real data. Such a comparison makes sense because *CellRanger* has been used for mapping/quantification of the 10X scRNA-seq data in thousands of studies, and the results have been validated by orthogonal experiments on many occasions. In these comparisons, the differences between tools arise not only from the differences in read mapping and read-to-gene assignment, but also from the differences in CB/UMI processing.

**Figure 4:**
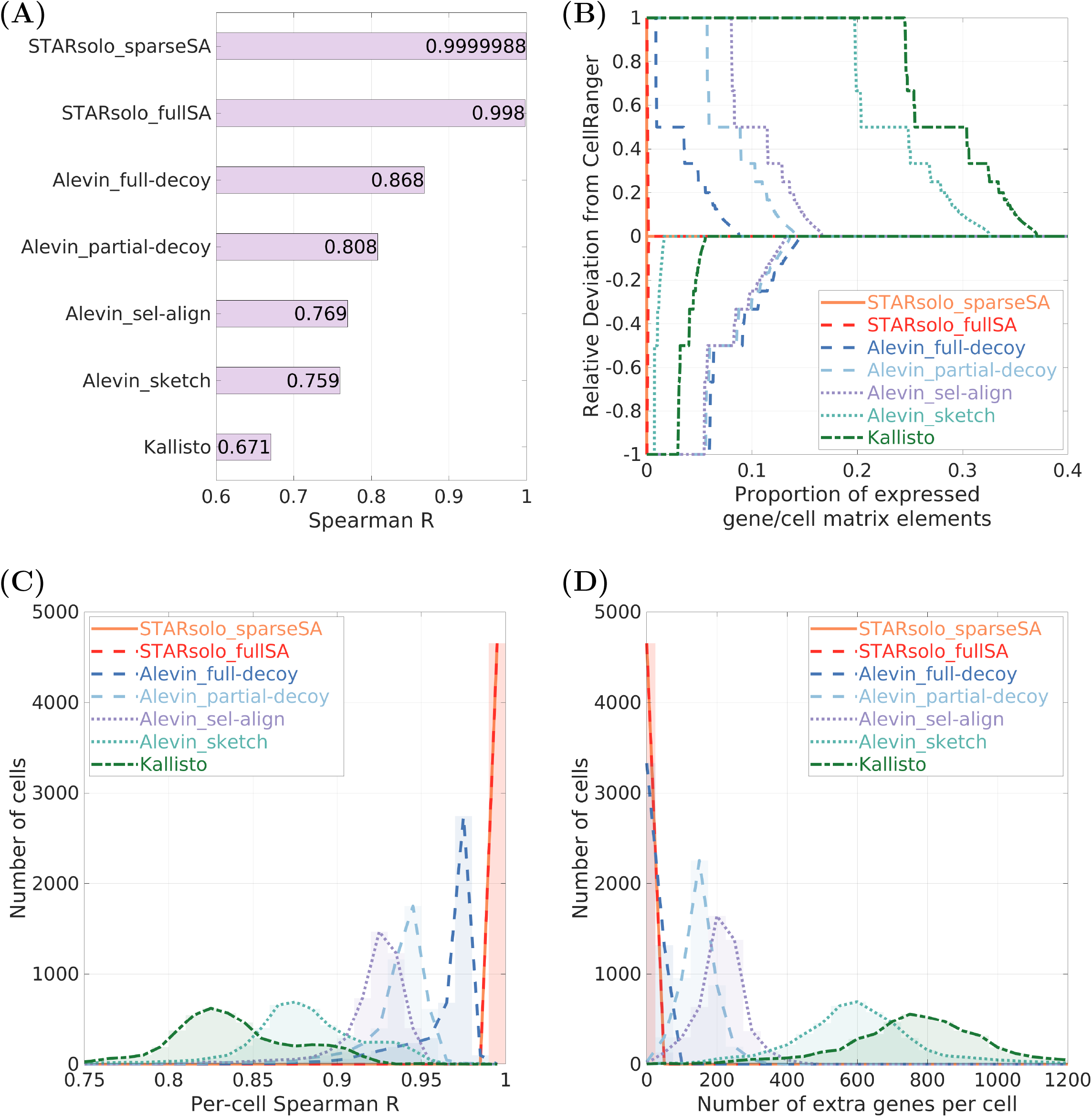
Mapping/quantification comparison for the **real** dataset 10X-pbmc-5k (Results 2.5). **(A):** Spearman correlation coefficient R vs *CellRanger* for gene/cell matrix elements. **(B):** Relative Deviation for gene/cell matrix elements with respect to the *CellRanger* counts. **(C):** Histogram of per-cell Spearman correlation coefficients vs *CellRanger*. **(D):** Histogram of the number of extra genes that were *not* detected by *CellRanger*, but were detected by each tool.

As *STARsolo* was run with parameters designed to match *CellRanger*’s gene quantification, we see that *STARsolo_sparseSA* (with sparse suffix array) results are nearly identical to *CellRanger*’s (which also uses sparse suffix array). In this example, out of ~17*M* non-zero gene/cell UMI counts, we only find a difference by one count in 85 gene/cell matrix elements (~0.0005%). *STARsolo_fullSA*, which utilizes a full suffix array, is still nearly perfectly correlated with *CellRanger*. The advantage of the full suffix array is in higher mapping speed, though it comes at the cost of higher RAM consumption (see Benchmarking section 2.7).

Other tools exhibit much lower agreement with *CellRanger*’s results. Similarly to the trends observed in the simulated data, pseudoalignment-to-transcriptome tools show the largest deviation from *CellRanger*. The agreement steadily improves for *Alevin* as it moves towards a more complete alignment to a fuller genome. In concordance with simulation results, *Alevin_sketch* and *Kallisto* predict expression for a large number of extra genes in each cell (~600 and ~800 on average) that are not expressed according to *CellRanger*, which constitutes a substantial proportion of all genes expressed in a cell (on average ~21% and ~26% respectively). We note that significant differences between *STARsolo*/*CellRanger* and *Kallisto* gene quantifications have been previously reported. [29, 30].

### 2.6 Differential gene expression is strongly influenced by mapping/quantification tools

To assess the effect of mapping/quantification algorithms on downstream analyses, we performed clustering and differential gene expression (DGE) calculations for the 10X-pbmc-5k dataset. We used *Scanpy* [31] and followed a standard pipeline (Methods 5.3) described for a similar dataset in [32–34]. As in the previous section, we used a common set of cells for all tools. Additionally, the clustering and cell type annotation were performed only for *CellRanger* (Figure S5) and transferred to all other tools.

We identified significantly differentially expressed marker genes for each cell type using the Wilcoxon rank-sum method and Benjamini-Hochberg multiple testing correction, requiring *p*_*adj*_<0.01. Figure 5A compares the sets of significantly overexpressed marker genes in each cluster between all tools and *CellRanger*. We observe a nearly perfect agreement for *STARsolo_sparseSA*/*fullSA*, and a decent agreement for *Alevin_full-decoy*, while *Kallisto* and *Alevin_sketch* deviate significantly from *Cell-Ranger*.

**Figure 5:**
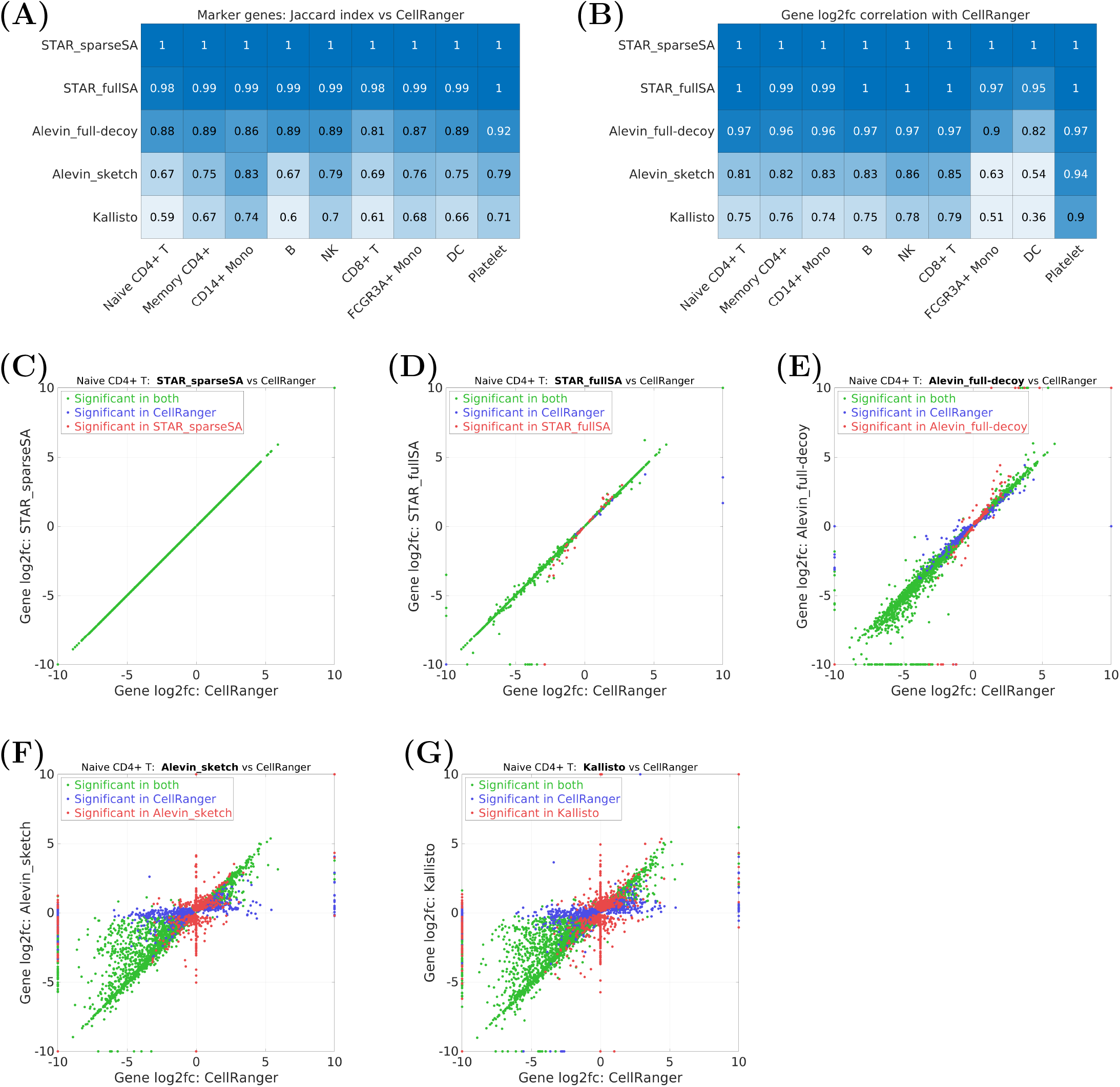
Differential gene expression in the **real** 10X-pbmc-5k dataset (Results 2.6). **(A):** Jaccard index for significantly overexpressed cluster marker genes, each tool vs *CellRanger*. **(B):** Pearson correlation between each tool and *CellRanger* for log2-fold-changes of significantly differentially expressed genes in each cluster. **(C)-(G):** Log2-fold-changes for significantly (*p*_*adj*_ < 0.01) differentially expressed genes in the *Naive CD4+ T* cluster, each tool vs *CellRanger*. The *log*_2_(*FC*) values were truncated at −10 and 10. The genes that were not detected by a tool were assigned *log*_2_(*FC*) = 0.

In Figures 5C-G we compare the log2-fold-changes (i.e. the effect sizes) for the significant marker genes of the largest cluster (*Naive CD4+ T cells*) for each tool vs *CellRanger* (see Supplementary Figures S6-S13 for other clusters). The Pearson correlation coefficients between these values are shown in Figure 5B.

In accord with our previous observations, pseudoalignment-to-transcriptome tools show poor agreement with *CellRanger*, which is especially pronounced for some of the smaller cell clusters (*FCGRA3+ Mono* and *Dendritic Cells*, Figures S11,S12). The red circles with *CellRanger*’s log_2_(*FC*)=0 on these plots represent the genes that were not detected by *CellRanger* in any cells, but were deemed significant by *Kallisto* and *Alevin_sketch* (likely False Positives). As before, we observe a close to perfect correlation with *CellRanger* for *STARsolo_sparseSA*/*fullSA*, and a generally good correlation for *Alevin_full-decoy*.

### 2.7 Benchmarks: trading accuracy for speed

To benchmark speed and Random Access Memory (RAM) usage, we ran all tools on the 10X-pbmc-5k dataset (Methods 5.4). Figure 6A shows the run-time, scaled to 100*M* reads, as a function of the number of threads. Good thread scalability is essential for taking full advantage of the current trend in processor technologies which increases the number of cores with each generation. We see that for most tools the processing speed saturates at ~16-20 threads, with the exception of *Kallisto*, whose performance saturates at ~8 threads.

**Figure 6:**
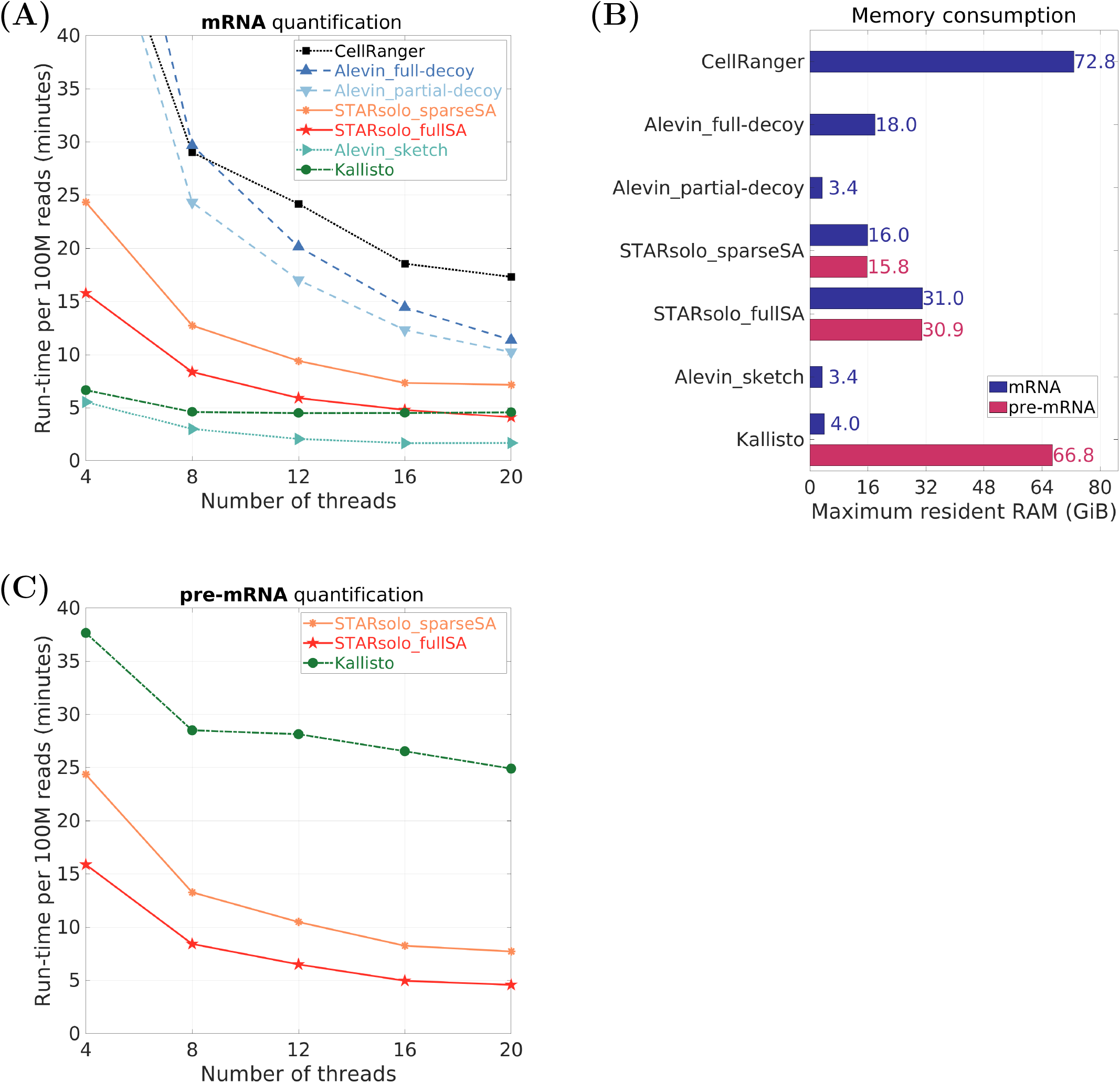
Speed and RAM benchmarks on the **real** 10X-pbmc-5k dataset (Results 2.7). **(A):** Run-time vs number of threads: wall-clock time scaled to 100 million reads, for mRNA quantification. **(B):** Maximum resident memory for mRNA (single-cell) and pre-mRNA (single-nucleus) quantifications, for runs with 20 threads. **(C):** Run-time vs number of threads for pre-mRNA quantification.

Pseudoalignment-to-transcriptome tools trade accuracy for high speed and low RAM usage. *Alevin_sketch* is ~2-3 times faster than *STARsolo_fullSA*. *Kallisto* is 2 times faster than *STARsolo_fullSA* with 8 threads, though this advantage diminishes as the number of threads increases and disappears at ~16 threads. On the other hand, among the highly accurate tools, *STARsolo* is the fastest: *STARsolo_fullSA* is 4.4 times faster than *CellRanger* and 2.8 times faster than *Alevin_full-decoy*.

Since transcriptome sequence length is ≲10% of the whole genome size, the pseudoalignment-to-transcriptome tools require a small amount of RAM (≲4 GiB, Figure 6B). On the other hand, for tools that map reads to the whole genome, RAM usage depends on the full genome size: for the human genome, *STARsolo_fullSA* uses 32 GiB. *STARsolo_sparseSA* trades speed for lower RAM usage: it uses 16 GiB of RAM, but it is 1.7 times slower than *STARsolo_fullSA*. While *CellRanger* also uses the sparse suffix array, its memory usage scales with the number of threads, taking 14.6 GiB of RAM for every 4 threads, requiring 73 GiB of RAM for 20 threads. *Alevin partial-decoy* uses a small part of the genome as a decoy, making its RAM consumption (3.4 GiB) as small as *Alevin_sketch*’s, and much smaller than *Alevin_full-decoy*’s (18 GiB).

All the comparisons and benchmarks up to this point were done for spliced (mature) mRNAs. We also performed benchmarks for another type of quantification: expression of pre-mRNAs. Such quantification is necessary for single-nucleus RNA-seq data which is widely used in cases where cell isolation is problematic (e.g. for brain cells). The majority of RNAs in a nucleus are unspliced pre-mRNAs, and thus quantifying snRNA-seq data requires counting reads that map to both exonic and intronic sequences. This is straightforward for *STARsolo*, since it maps reads to the whole genome, and only the read-to-gene assignment algorithm has to be adjusted. On the other hand, the pseudoalignment-to-transcriptome tools require significant modifications of their algorithms to include intronic sequences, resulting in many-fold speed reduction and RAM usage increase.

Figures 6B,C show that *STARsolo*’s speed and RAM usage stay the same for pre-mRNA quantification, while *Kallisto*’s speed is reduced by a factor of ~5, and its RAM usage increases by a factor of ~17 to a total 68 GiB. Thus, for single-nucleus pre-mRNA quantification, both *STARsolo_sparseSA* and *STARsolo_fullSA* are up to ~2 and ~5 times faster than *Kallisto*, while consuming significantly less RAM.

### 2.8 Beyond gene expression: cell-type specific alternative splicing

Similarly to bulk RNA-seq, single-cell data contain information about other transcriptomic features, such as splicing, isoform expression, polyadenylation, allele-specific expression, etc. Characterizing these transcriptomic features in a cell-type-specific manner may enable discovery of interesting biological phenomena beyond gene expression. However, most analyses of droplet-based scRNA-seq data are focused on gene expression, with a few notable exceptions: [35–40].

It is a common preconception that the 3′-end cloning bias prevents studying splicing in droplet-based scRNA-seq data. Here we illustrate *STARsolo* capabilities for detecting alternative splicing in scRNA-seq data by analyzing the data from the Tabula Muris [41] project. Tabula Muris contains 10X scRNA-seq libraries for ~100 thousand cells from 20 *Mus musculus* organs and tissues. We found that a large proportion (~20-40%) of the reads in these datasets are spliced, allowing quantification of splice junctions (SJ) in different cell types.

We used gene/cell and SJ/cell counts output by *STARsolo* to calculate SJ usage (*U*_*sj*_) in each cell-type using the pseudobulk approach (see Methods 5.5). For each SJ and cluster, the SJ usage metric is defined as total SJ expression in the cluster normalized by the expression of the corresponding gene. The clustering and cell type annotations were generated by the Tabula Muris project based on gene expression. To identify significant inter-cluster SJ usage switching, we use Wilcoxon rank-sum test for bootstrapped cell samples from each cluster, and Benjamini-Hochberg multiple testing correction.

In Figures 7A,B we show the differential SJ usage between a pair of cell clusters. As expected, for most of the SJs, the usage changed only slightly between the cell types. Still, there are many SJs whose usage considerably changes between the clusters, resulting in cell-type dependent differential expression of the corresponding alternatively spliced isoforms. Figures 7C-F show the number SJs with significant cell-type specific SJ usage for several tissues, aggregated over cell type pairs in each tissue, as well as aggregated over all tissues. We see that hundreds of SJs undergo substantial usage switches in each tissue. Of course, because of low sequencing depth and 3′-end cloning bias, only a small portion of significant cell-type specific SJs can be detected in droplet-based scRNA-seq protocols. However, we believe that such analyses may lead to discovery of some interesting examples of biologically important cell-type specific alternative splicing.

**Figure 7:**
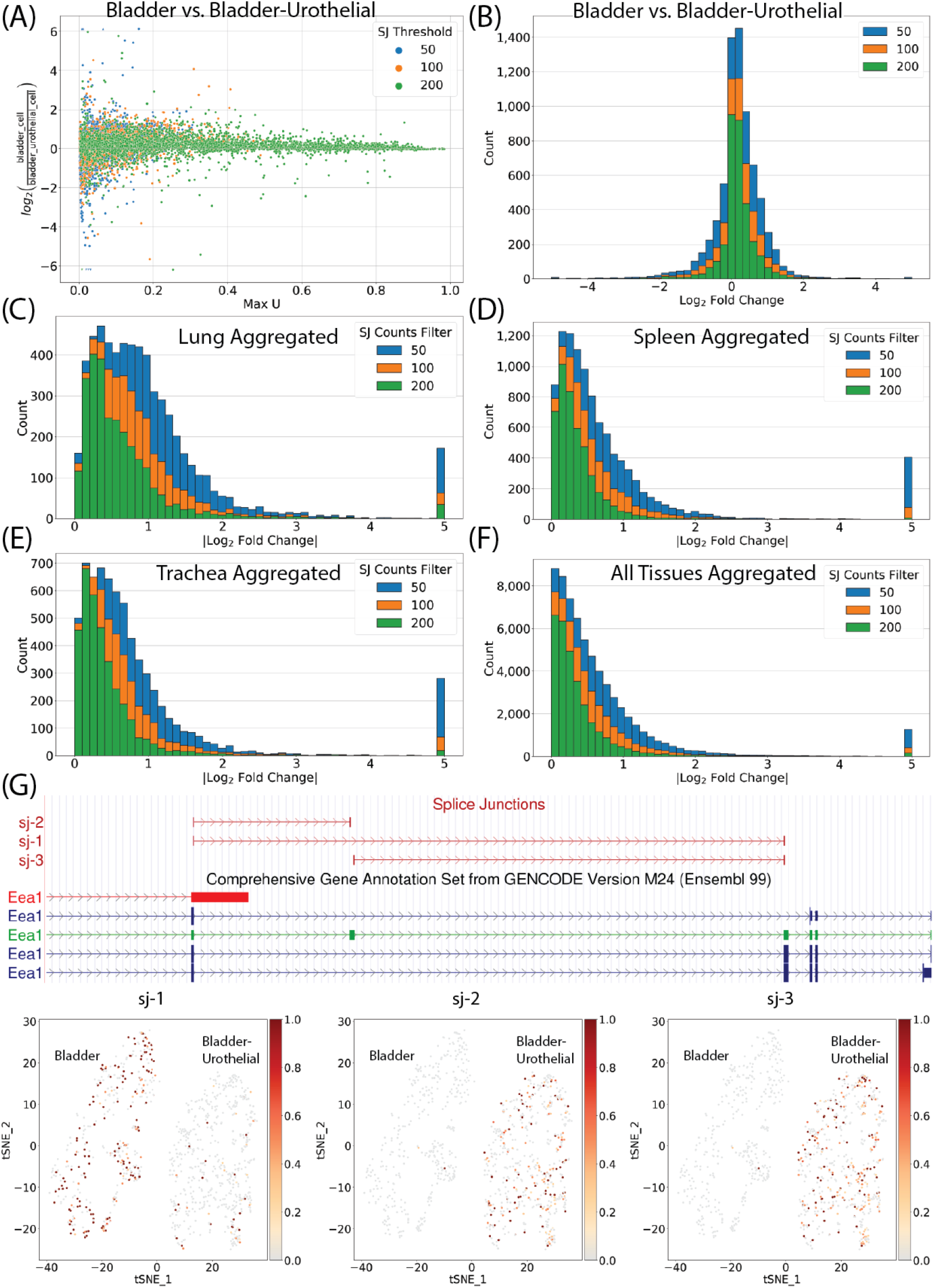
Cell-type-specific alternative splicing in the Tabula Muris dataset (Results 2.8). **(A):** SJ usage (*U*_*sj*_) fold change vs *U*_*sj*_ maximum in Bladder vs Bladder-Urothelial cell types. Only SJs with significant usage switching (*p*_*adj*_ < 0.05) are shown. SJ thresholds limit mark different minimum SJ counts (50, 200, 100) in one of the clusters. **(B):** Histogram of *U*_*sj*_ log2-fold-changes in Bladder vs Bladder-Urothelial cell types. **(C,D,E):** Histograms of *U*_*sj*_ log2-fold-changes aggregated for all cell-type pairs in Lung, Spleen, Trachea. **(F):** Histograms of *U*_*sj*_ log2-fold-changes aggregated for all cell-type pairs in all Tabula Muris tissues. **(G):** Example of an exon-skipping event in Bladder vs Bladder-Urothelial clusters: browser snapshot and UMAP-embedded *U*_*sj*_ usage values.

## 3 Discussion

In this work we presented *STARsolo*, an accurate, fast and versatile tool for mapping and quantification of single-cell RNA-seq data

To assess *STARsolo*’s accuracy and compare it with other mapping/quantification tools, we developed a method for simulating scRNA-seq reads, which closely resembles real data. Simulations play an important role in assessing quality of the results, as they establish the ground truth not available for real data. We believe that our simulation procedure will be helpful in guiding future improvements of mapping/quantification algorithms.

Using simulated data, we demonstrated the high accuracy of *STARsolo*’s gene quantification. On the other hand, our simulations show a significantly lower accuracy for pseudoalignment-to-transcriptome tools (*Kallisto* and *Alevin_sketch*). This reduction in quantification accuracy is caused by intronic reads, which constitute a large proportion of scRNA-seq libraries. The pseudoalignment-to-transcriptome algorithms force intronic reads to map to spurious genes, resulting in hundreds false positive genes in each cell. We note that the developers of *Salmon/Alevin* addressed this problem by moving away from the pseudoalignment-to-transcriptome to a more accurate selective alignment to both transcriptome and genome decoy [28]. As a result, *Alevin_full-decoy* achieves high (as good as *STARsolo*) agreement with the simulated truth.

Our benchmarking of processing speed and RAM usage showed strong trade-offs between accuracy and speed/RAM usage. The least accurate pseudoalignment-to-transcriptome tools are also the fastest and consume the least amounts of RAM. On the other hand, among the highly accurate tools, *STARsolo* with full suffix array is 4.4 times faster than *CellRanger* and 2.8 times faster than *Alevin_full-decoy*. *STARsolo* can use a sparse suffix array, reducing RAM consumption 2-fold (from 32 to 16 GiB for human genome) at the cost of 1.7-fold reduction in speed, which is still faster than *CellRanger* or *Alevin_full-decoy*.

Furthermore, *STARsolo*’s RAM usage and speed do not depend on the type of transcript/gene annotations used to assign reads to genes. Contrarily, the computational efficiency of the pseudoalignment-to-transcriptome tools drastically deteriorates when intronic reads have to be counted in addition to exonic reads. This is required, for instance, for quantifying pre-mRNA in single-nucleus RNA-seq data, or unspliced transcripts in RNA Velocity calculations[13]. We showed that for pre-mRNA quantification, *STARsolo* is up to 5 times faster than *Kallisto* and consumes less RAM (32 GiB vs 68 GiB).

For real scRNA-seq data, we compared *STARsolo* and other tools with *Cell-Ranger*. We demonstrated that *STARsolo* with proper parameters can match *Cell-Ranger*’s results almost perfectly, while other tools show significant differences. While *CellRanger*’s quantification is not the ground truth (which is not available for real data), we still believe it is an important reference point, since *CellRanger* has been widely used for analyzing thousands of scRNA-seq datasets generated with 10X Genomics technology. Many of the biological discoveries in these studies were validated with alternative experimental techniques, thus confirming the overall validity of the *CellRanger* algorithms.

An additional advantage of replicating *CellRanger* results is that it simplifies comparing a new experiment with existing data. *STARsolo* needs to be run only on the new dataset, and its output can be directly compared to the old *CellRanger* quantifications.

To replicate *CellRanger* results we carefully dissected *CellRanger* procedures for read trimming, read-to-gene assignment, cell barcode demultiplexing, UMI deduplication and cell filtering, in some cases using the source code, available for the older *CellRanger* versions, while in other cases reverse engineering the algorithms from the observed output. This signifies that we validated *CellRanger*’s approaches *in silico*. We believe this is an important service to the research community, as the *CellRanger* is widely used, but its methodology has not been well documented or peer-reviewed.

In addition to significantly higher speed and lower RAM usage, *STARsolo* has other advantages over *CellRanger*. *STARsolo* can take into account multi-gene reads, which are presently disregarded by the *CellRanger*. Including multi-gene reads is important for detecting certain classes of biologically important genes (e.g. paralogs). *STARsolo* has several different strategies for taking into account the multi-gene reads, which we tested on simulations with multi-gene reads. We observed that multi-gene quantification options increase the sensitivity, “rescuing” the multi-gene reads and thus detecting genes that are supported by multi-gene reads only. However, at the same time, some of the multi-gene reads are not properly resolved, which slightly decreases the precision.

Another advantage of *STARsolo* is its ability to process data from multiple scRNA-seq platforms, while *CellRanger* is locked into 10X Genomics products, by both software design and licensing [42]. A multitude of scRNA-seq methods have been published, with a variety of read barcoding schemes. To accomodate a large range of single cell technologies, *STARsolo* uses a flexible system to describe CB/UMI geometry, suppporting multiple CBs with distinct passlists, variable CB lengths, and anchored to a specified adapter sequence.

Yet another important feature of *STARsolo* is its ability to quantify transcriptomic features beyond gene expression, such as splice junctions and spliced/unspliced transcripts, required for RNA Velocity calculations [13]. By analyzing the data from multiple tissues in the Tabula Muris dataset, we showed that despite strong 3′-end bias, it is possible to detect cell-type-specific splice junction usage in droplet-based scRNA-seq. *STARsolo* can also output a standard BAM file containing read alignments and error-corrected CBs and UMIs, which can be used for a variety of downstream analyses, such as differential splicing [36], alternative polyadenylation [39], allele specific expression [38, 40], fusion detection, etc.

Last, but not least, *STARsolo* is a truly open source software, distributed under the unrestrictive MIT license on GitHub. The entire history of *STAR* releases is available, allowing for a precise reproducibility of older results. Importantly, we welcome contributions to the *STAR* code, as well as comments, suggestions and feature requests.

There are multiple areas where *STARsolo* performance can be enhanced. We are working on improving quantification for multi-gene reads, adapting additional single-cell technologies (e.g. CITE-seq-like [43] epitope indexing and scATAC-seq [44, 45]), quantifying other transcriptomic features, as well improving documentation and creating tutorials. We believe that owing to its uncompromised accuracy, robust computational efficiency and technological versatility, *STARsolo* will become a useful tool for single-cell genomic studies.

## 4 Acknowledgments

We would like to thanks ENCODE RNA working group for fruitful discussions. This work was supported by the National Human Genome Research Institute of the National Institutes of Health under award number R01HG009318. The content is solely the responsibility of the authors and does not necessarily represent the official views of the National Institutes of Health.

## 5 Methods

### 5.1 *STARsolo* implementation

*STARsolo* is built directly into the RNA-seq aligner *STAR* [12], and can be run similarly to standard *STAR* bulk RNA-seq alignment, specifying additionally the single-cell parameters such as barcode geometry and passlist (a.k.a. whitelist). Users can adjust *STAR* mapping parameters at will, allowing for sample and task specific customization. Multiple options are also available for tweaking cell barcode demultiplexing and UMI deduplication. One of the design goals was to closely match the gene/cell count matrix values and formatting of the 10X *CellRanger*, a well established tool used in thousands of publications for mapping/quantification of 10X scRNA-seq data.

#### 5.1.1 Workflow

In the mapping step, *STARsolo* inputs reads from FASTQ files, or, optionally, from unmapped or mapped BAM files (the latter is useful for reprocessing *CellRanger* BAM files). The reads are aligned to the reference genome, and alignments are checked for concordance with the transcript models to assign them to genes (5.1.2) and/or other features (5.1.3). Simultaneously, if the barcode passlist is provided, the cell barcodes are matched against the passlist, allowing for up to one mismatch (5.1.4). In addition to simple barcoding schemes, *STARsolo* has a flexible mechanism (5.1.5) for supporting complex barcode schemes. For each read, its CB, UMI and gene are recorded into temporary files. It is possible to save the temporary files for postprocessing with a different pipeline, and in the future we are planning to implement standardized formatting similar to the BUS and RAD formats implemented in *Kallisto-bustools* [9] and *Alevin-fry* [10].

In the quantification step, *STARsolo* reads the CB, UMI and gene information from temporary files, and performs CB demultiplexing and UMI deduplication (5.1.6). The deduplicated UMIs are counted towards their gene/cells, multi-gene reads are recovered (5.1.7), and the raw gene/cell count matrix is generated. Finally, the cell calling (filtering) is performed to eliminate cell barcodes containing ambient RNA (5.1.8), and the filtered gene/cell count matrix is generated.

#### 5.1.2 Read mapping and assignment to genes

*STARsolo* aligns reads to the whole genome, it uses the same genome index as *STAR*. In principle, it is possible to align reads only to the transcriptome, by generating the index for transcript sequences, which will drastically (~10-fold) reduce the memory requirements. However, because of high content of intronic reads in scRNA-seq libraries, aligning scRNA-seq reads to transcriptome only results in severe accuracy reduction (see Results 2.3).

For each read alignment, we find a set of concordant annotated transcripts. The strandedness of the scRNA-seq library is user-specified with the --soloStrand option. An alignment is considered concordant with a transcript if all of its alignment blocks are contained within transcript exons. If splice junctions are present in the alignment, they have to agree with the transcript junctions. From the set of concordant transcripts we create a set of corresponding concordant genes. If this gene set contains only one gene, the read is considered unique-gene. Note that reads mapping to multiple loci in the genome will be considered unique-gene if only one of the alignments is concordant with one gene, or if all the alignments are concordant with the same gene. These rules follow *CellRanger*’s read-to-gene assignment policy.

For multi-gene reads, whose concordant gene set contains two or more genes, the read-to-gene(s) assignment is performed probabilistically at a later stage (5.1.7).

#### 5.1.3 Quantifying other transcriptomic features

In addition to quantifying gene expression, *STARsolo* can quantify other transcriptomic features. All the features (including the standard --soloFeatures *Gene*) can be quantified simultaneously in a single *STARsolo* run.

--soloFeatures *GeneFull* produces per-cell counts of reads that overlap the entire gene locus, i.e. both exons and introns. It will count reads originating from both mature mRNA and pre-mRNA, and hence it is appropriate for quantifying gene expression in single-nucleus RNA-seq experiments.
--soloFeatures *SJ* quantifies splice junctions by calculating per-cell counts of reads that are spliced across junctions. It will count spliced reads across annotated and unannotated junctions, thus allowing analysis of inter-cell alternative splicing (see Results 2.8) and detection of novel splice isoforms.
--soloFeatures *Velocyto* performs separate counting for spliced, unspliced and ambiguous reads, similar to the *Velocyto* tool [13]. Its output can be used in the RNA-velocity analyses to dissect the transcriptional dynamics of the cells.

#### 5.1.4 CB passlisting and error correction

*STARsolo* converts CB sequences into a 2-bit binary representation, allowing up to 32-mer CBs that are stored in 8 bytes each. *STARsolo* uses a simple binary search algorithm to exactly match the CB to the barcode passlist (--soloCBmatchWLtype *Exact*).

--soloCBmatchWLtype *1MM* allows for a Hamming distance of ≤1 (i.e. up to 1 mismatch) when matching CBs to the passlist. This is done by creating all possible 1-base substitutions of the original barcode sequence and checking them against the passlist with the binary search. The CBs that map to multiple passlist barcodes with different mismatches are discarded. While this is not the most efficient algorithm, it only marginally increases overall run-time.
--soloCBmatchWLtype *1MM multi\** options allow to recover CBs that map to multiple passlist barcodes with different mismatches, following the procedure introduced by *CellRanger*, that estimates the posterior probability for each passlist match, based on the sequencing quality score and the number of reads exactly matching each passlist barcode.

For scRNA-seq technologies that do not provide a barcode passlist, *STARsolo* can operate without the passlist, outputting all CBs without error correction. If CB error correction is desired, users can perform passlisting based on the uncorrected results of the 1st pass, and then provide the passlist to the 2nd pass of *STARsolo*.

#### 5.1.5 Complex Cell Barcodes

To support technologies that use non-trivial barcode geometry (e.g. inDrop [46]), we implemented a flexible scheme for specifying CB and UMI positioning. The CB and UMI positions are specified with `--soloCBposition` and `--soloUMIposition` with respect to either barcode read start/end or adapter sequence (given in `--soloAdapterSequence`) start/end. Multiple CB positions can be specified with corresponding barcode passlists. Varying CB lengths are allowed in each passlist. This highly adaptable barcoding scheme allows processing a wide range of currently available scRNA-seq protocols.

#### 5.1.6 UMI deduplication and gene/cell UMI counts

Similarly to CB error correction, most scRNA-seq tools perform UMI deduplication with error correction. For instance, *Alevin* [7] implements a sophisticated UMI deduplication algorithm based on counting monochromatic arborescences of parsimonious UMI graphs, while recently released *Alevin-fry* [10] additionally implements several simpler methods. Contrarily, *Kallisto* does not perform UMI error correction. We implemented multiple options for various flavors of UMI error correction in *STARsolo*:

--soloUMIdedup *Exact* performs UMI deduplication without error correction, collapsing UMIs with identical sequences.
--soloUMIdedup *1MM All* performs 1-mismatch error correction, collapsing all UMIs within a Hamming distance of 1 from each other. This is equivalent to finding the number of connected components in a graph whose nodes are UMIs, and edges represent Hamming distance 1.
--soloUMIdedup *1MM Directional\** options use the “directional” approach implemented in *UMItools*, which collapses two UMIs with Hamming distance =1 only if the read number for the “winning” UMI is sufficiently higher than for the other UMI.
--soloUMIdedup *1MM CR* options closely mimics the *CellRanger* algorithm. It first sorts the UMI by the number of reads per UMI, and then alphabetically. Next, it moves through the sorted list, collapsing UMIs with Hamming distance ≤ 1.

While the multi-gene reads are not considered in this step, we can still encounter a UMI that maps to multiple genes. For such “gene-inconsistent” UMI, different reads map uniquely to different genes, which should not be possible under the assumption that each UMI originates from one RNA molecule. *STARsolo* has several options to filter out such UMIs:

--soloUMIfiltering *MultiGeneUMI* compares the number of reads for each gene for a given UMI, and keeps only the gene with the maximum number of reads. If two or more genes contain the same largest number of reads, the UMI is discarded.
--soloUMIfiltering *MultiGeneUMI CR* is similar to the previous option, but matches more closely *CellRanger*’s filtering, which is performed both before and after UMI error correction.
--soloUMIfiltering *MultiGeneUMI All* is the most stringent option, and filters out all UMIs that contain unique-gene reads mapping to different genes.

The gene/cell count matrix for unique-gene reads/UMIs is generated by simply counting the number of collapsed (deduplicated) UMIs for each gene/cell. In the current *STARsolo* implementation, the quantification step is performed after CB/UMI/gene information for all reads is loaded into RAM. This, in principle, is not as memory-efficient as the approach first implemented in *Kallisto-bustools* and later in *Alevin-fry*, which first sort the reads by CB, allowing CB/UMI processing in small batches. In practice, *STARsolo*’s RAM usage is dominated by the whole genome suffix array: 16 GiB for sparse and 32 GiB for full human suffix arrays. Suffix arrays are unloaded from RAM before the quantification step, which utilizes 16 bytes per read to record CB/UMI/gene information (= 8 + 4 + 4 bytes). Thus the maximum RAM consumption will not exceed the suffix array size unless the sample contains more than 1 Billion mapped reads for *STARsolo_sparseSA* and 2 Billion reads for *STARsolo_fullSA*.

#### 5.1.7 Multi-gene reads

Multi-gene reads are concordant with (i.e. align equally well to) transcripts of two or more genes. One class of multi-gene read are those that map uniquely to a genomic region where two or more genes overlap. Another class are those reads that map to multiple loci in the genome, with each locus annotated to a different gene.

Including multi-gene reads allows for more accurate gene quantification and, more importantly, enables detection of gene expression from certain classes of genes that are supported only by multi-gene reads, such as overlapping genes and highly similar paralog families.

First, we collect all multi-gene reads with the same CB and UMI. Since they originate from the same RNA molecule, we compute the intersection of gene sets from these reads, creating a multi-gene UMI gene set. Next, the multi-gene UMI is assigned probabilistically to one or more genes from its gene set, using one of the following options:

--soloMultiMappers *Uniform* uniformly distributes the multi-gene UMIs to all genes in its gene set. Each gene gets a fractional count of 1/*N*_*genes*_, where *N*_*genes*_ is the number of genes in the set. This is the simplest possible option, and it offers higher sensitivity for gene detection at the expense of lower precision.
--soloMultiMappers *EM* uses Maximum Likelihood Estimation (MLE) to distribute multi-gene UMIs among their genes, taking into account other UMIs (both unique- and multi-gene) from the same cell (i.e. with the same CB). Expectation-Maximization (EM) algorithm is used to find the gene expression values that maximize the likelihood function. Recovering multi-gene reads via MLE-EM model was previously used to quantify transposable elements in bulk RNA-seq [18] and in scRNA-seq [7, 8].
--soloMultiMappers *Rescue* distributes multi-gene UMIs to their gene set proportionally to the sum of the number of unique-gene UMIs and uniformly distributed multi-gene UMIs in each gene [15]. It can be thought of as the first step of the EM algorithm.

#### 5.1.8 Cell calling (filtering)

The outcome of the previously described steps is the *raw* gene/CB count matrix. It typically contains hundreds of thousands of CBs, while only a few thousand cell have been provided as input to the assay. The cell calling step aims to filter out the CBs that correspond to empty droplets, i.e. contain ambient RNA rather than true cells. Multiple methods have been developed to perform the cell filtering [4, 7, 47], and these tools can be directly applied to the raw count matrix generated by *STARsolo*.

To achieve agreement with *CellRanger* for the filtered gene/cell count matrix, we implemented the following cell calling options:

--soloCellFilter *CellRanger2.2* was used by the older versions of *CellRanger* and is based solely on the total number of UMIs per CB [14]. It filters out cells whose UMI count is <10% of the robust maximum count, which is defined as the 99% of UMI counts in all detected barcodes. This approach is somewhat similar to the *knee-point* thresholding used in [48]. While this approach works well for homogeneous cell populations, it fails to recover the cells with low RNA content in heterogeneous samples (e.g. tumor microenvironment samples that contain a mixture of large tissue cells and small immune cells).
--soloCellFilter *EmptyDrops CR* replicates the cell calling in newer *CellRanger* versions, which is based on the *EmptyDrops* approach [47]. *EmptyDrops* recovers small cells from complex samples by detecting CBs whose gene expression significantly deviates from the ambient RNA profile. *CellRanger* implementation differs from the *EmptyDrops* algorithm in several aspects, thus producing moderately different results. In particular, it uses the multinomial distribution as described in the first version of the *EmptyDrops* [49] rather than Dirichlet-multinomial distribution [47]. In addition, *CellRanger* makes different choices with respect to the choice of cells with high UMI counts that do not undergo filtering, the choice of candidate low-count cells to be tested against ambient RNA profile, and the choice of CBs that are assumed to contain ambient RNA.

### 5.2 Simulating scRNA-seq reads

The main idea of our simulation is to get the information about CB/UMI and read position on a transcript (or genome) from the real scRNA-seq data. The advantage of this approach is that it does not require to build a generative statistical model of read distribution along the transcript length, as is done for the bulk quantification algorithms [50–52]. Building a generative model for scRNA-seq is a non-trivial task, because read distribution along the exons and introns is highly irregular, owing to non-uniformity and non-linearity of Reverse transcription polymerase chain reaction (RT-PCR).

Our simulation method consists of the following steps:

1. Select a real scRNA-seq datasets to be the basis of the simulations, the corresponding passlist, as well as genome asssembly and transcript/gene annotations.
2. Remove reads with CBs that do not exactly match any entries in the CB-passlist. Since only exact passlist matches are kept, our simulation does not test the CB error correction algorithms. Also remove UMIs with undefined (“N”) bases.
3. Map the real dataset to the *combined* full genome and transcriptome. For this task we use highly accurate short read aligner *BWA-MEM* [27] with -a option, which outputs *all* detected alignments. It is crucially important to map reads to transcriptome and genome sequences simultaneously, because a large proportion of reads in scRNA-seq data originate from non-exonic (mostly intronic) sequences that are absent from the transcriptome sequence.
4. For each read, we select those *BWA-MEM* alignments whose score is equal to the highest score for this read.
5. If read CB/UMI combination was not yet observed, randomly choose one of the top-scoring read alignments. If the top-scoring set contains alignments to both transcripts and genome, we choose only among transcript alignments. The selected alignment determines the origin of the CB/UMI-specified RNA molecule: it can be either one specific transcript, or a general genomic (non-transcript) locus.
6. If the CB/UMI is assigned to a transcript, add 1 to the corresponding gene/cell in the *true* count matrix.
7. If a read CB/UMI combination was already observed, choose the alignment with the same transcript/genomic locus as was recorded the first time it was observed. If the top-scoring set does not contain proper transcript/genomic locus, this read is discarded. This procedures ensures that each CB/UMI corresponds to only one transcript or genomic locus.
8. For the selected alignment, extract the “true” sequence from the transcript or genomic sequence. To simulate sequencing errors, make random sequence substitutions with a nominal error rate. In this study we used mismatch error rate of 0.5%, which is similar to the one observed in realistic Illumina sequencing.
9. Record the simulated sequence, as well as CB/UMI in read 1/2 FASTQ files, which, together with the true gene/cell count matrix, comprise the simulated dataset.

We note that all the properties of our simulated dataset are defined by the real scRNA-seq dataset (which served as the basis), genome assembly, transcript/gene annotations, and the mismatch error rate (the only adjustable parameter).

### 5.3 Real data: clustering and differential gene expression

We used a common set of 4655 cells (Results 2.5) as a starting point for the *scanpy 1.6.0* [31] pipeline described in [34], which, in turn is based on the *Seurat* tutorial [33]:

1. We further filtered the cells that contain < 200 genes or > 20% of mitochondrial reads based on *CellRanger* counts, resulting in the final set of 4, 473 cells which was used for all tools.
2. For each tool, genes detected in less than 3 cells were excluded.
3. Counts were normalized to 10000 reads per cell, a pseudocount of 1 was added and the natural log transformation was performed. These normalized expression values were used for differential gene expression calculations (see below).
4. For UMAP embedding, neighborhood graph calculation and clustering, the expression values were scaled to unit variance and zero mean across the cells and truncated at a maximum value of 10.
5. 3, 000 highly variable genes were selected using the ‘seurat v3’ algorithm.
6. The shared neighborhood graph was produced for 20 nominal nearest neighbors, based on euclidean distance for the top 20 PCA components
7. We clustered *CellRanger*’s neighborhood graph using the Leiden clustering algorithm, adjusting the resolution to obtain 9 clusters.
8. We assigned cell types to clusters using marker genes from [33], see Figure S5.
9. We transferred *CellRanger*’s cell-cluster labels obtained above to all other tools for differential gene expression comparisons.
10. The differentially expressed genes in each cluster for each tool were detected using *scanpy*’s implementation of the Wilcoxon rank-sum test, which was previously found to be one of the top performing methods [53].

### 5.4 Benchmarking speed and RAM consumption

All benchmarks were performed on the workstation with two Intel Xeon Gold 6140 processors (total 36 cores), 384 GiB of DDR4/2666 RAM, and 8 12TB SATA hard drives in RAID0 configuration.

The benchmarking parameters were the same as used for the real scRNA-seq data comparisons (Supp. Table 1). Output of the Linux command `/usr/bin/time -v` was parsed to extract run-time (“Elapsed (wall clock) time”) and maximum RAM consumption (“Maximum resident set size”). Since *CellRanger* spawns multiple processes, a custom made script was needed to track its total memory consumption. Each tool was run 5 times, and the mean run-time was calculated.

### 5.5 Alternative splicing analysis

For analysis of alternative splicing, we reprocessed the 10X libraries generated by the Tabula Muris consortium [41]. *STAR* genome index was generated for *GRCm38* mouse genome assembly with *Gencode M20* annotations. *STARsolo* was run with --soloFeatures *Gene SJ* to output both gene/cell and SJ/cell count matrices. Only annotated SJs and only reads that mapped uniquely to a single splice junction were considered in this analysis.

Our analysis utilizes the pseudobulk approach: the SJ expression values are averaged over cell types which are determined by clustering the cells using gene expression. We used the clustering and cell type annotations provided by Tabula Muris [41]. To calculate the average relative SJ usage in a cluster (denoted as U), we normalized SJ counts by the expression of the gene the SJ belongs to: we sum the SJ counts over the cells in the cluster and divide it by sum of the counts for the corresponding gene:

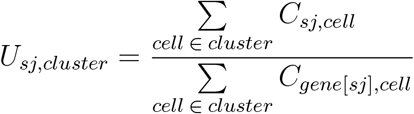

To estimate the variance of each SJ in a cluster, we used a bootstrapping approach. *U*_*sj,cluster*_ were calculated on 100 sets of cells randomly chosen with replacement from each cluster. Wilcoxon rank-sum test was applied to the bootstrapped *U*_*sj,cluster*_ values to calculate the *p*-values, which were adjusted for multiple testing using the Benjamini-Hochberg procedure. Median values of *U*_*sj,cluster*_ were used to represent average relative SJ usage in each cluster.

To avoid low-expressed genes, we only considered SJs that belonged to genes with expression greater than the 75th percentile in each cluster. Also, we only considered clusters with ≥200 cells. More detailed description of our splicing pipeline can be found in Supp. Methods 2.1.

### 5.6 Code availability

*STARsolo* is built into *STAR* and is available on: https://github.com/alexdobin/STAR. The scripts for the analyses presented in this manuscript are available on: https://github.com/dobinlab/STARsoloManuscript.

