## Supplementary Figures and Notes for "STARsolo: accurate, fast and versatile mapping/quantification of single-cell and single-nucleus RNA-seq data"

### Supplementary Materials

#### 1 Supplementary Figures

**Figure S1:** Simulations **without multi-gene reads** (Results 2.2).
**(A):** Matrix of Spearman correlation coefficients for gene/cell matrix elements.
**(B):** Mean per-cell Spearman correlation coefficient R; Mean Absolute Relative Deviation; Mean proportion of false positive genes per cell.

**(A)**

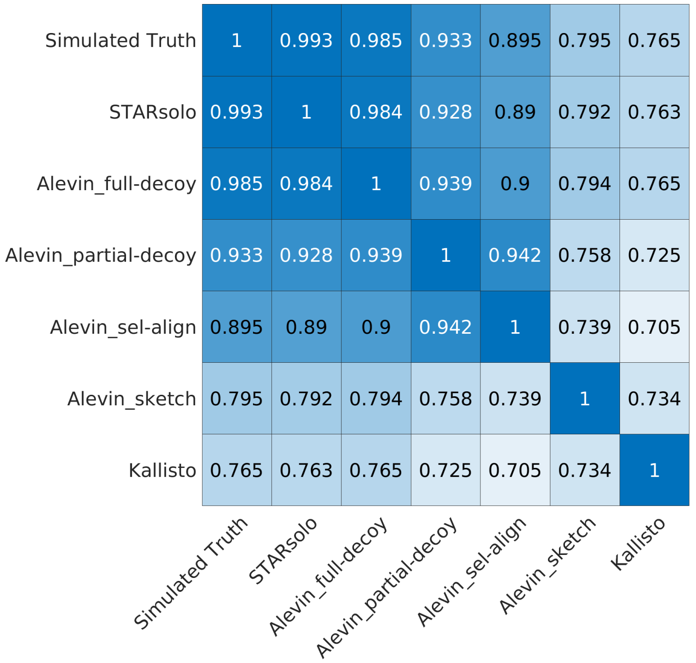


**(B)**

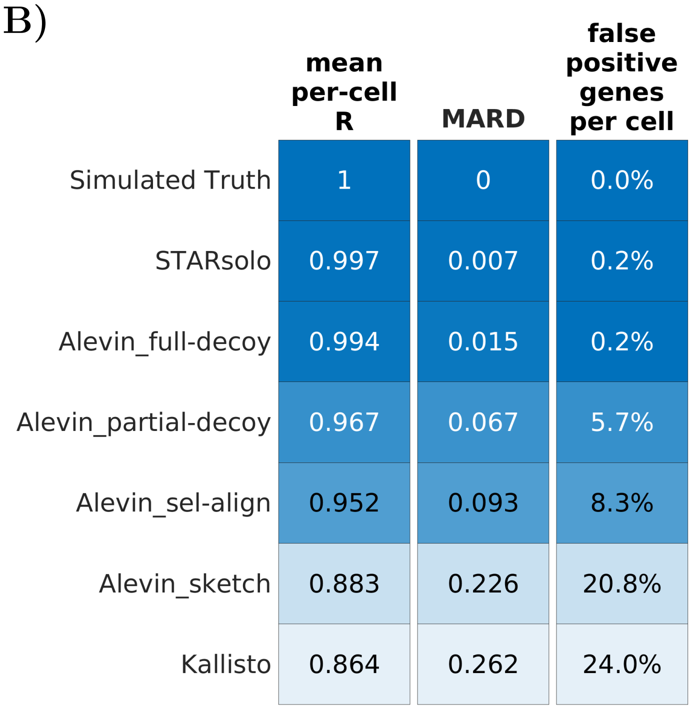

**Figure S2:** Simulated data **without multi-genic reads** and **without non-exonic reads**: Spearman correlation coefficient for gene/cell matrix elements between each tool and the truth (Results 2.2).

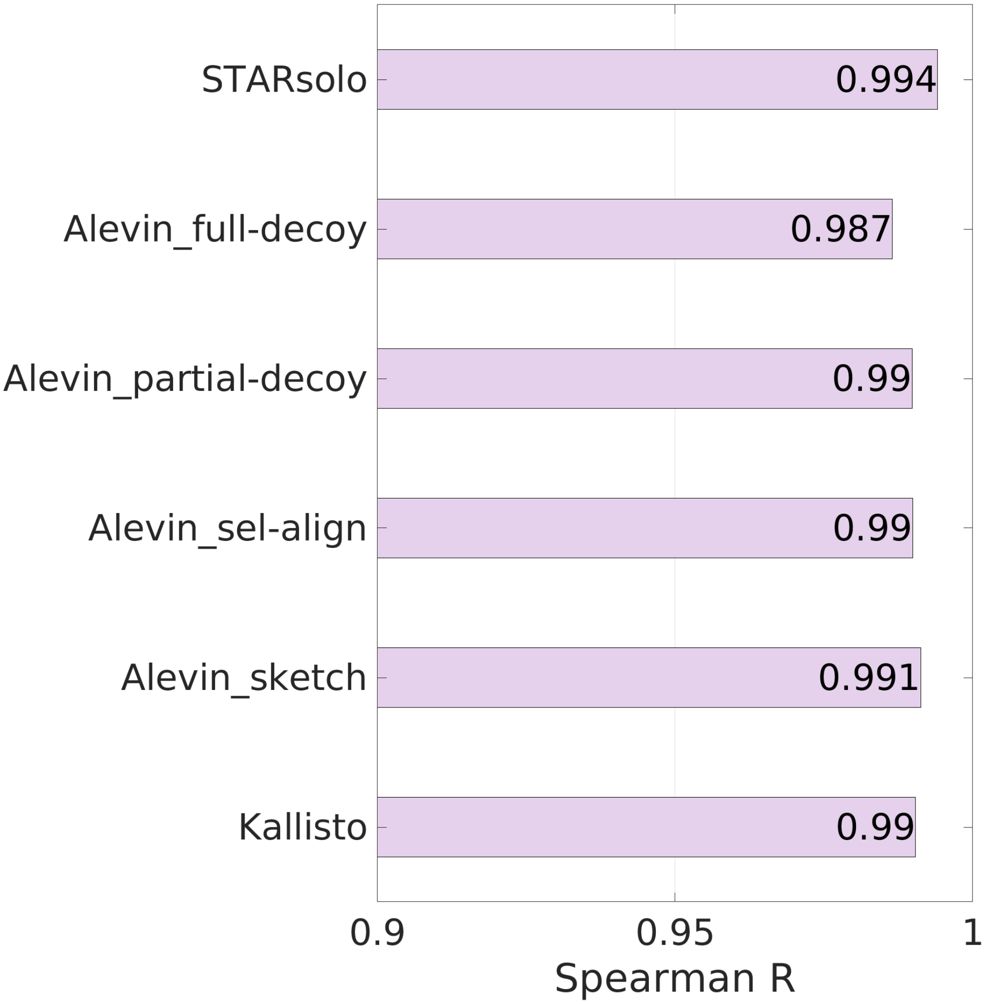

**Figure S3:** Simulations **with multi-gene reads** (Results 2.4).
**(A):** Matrix of Spearman correlation coefficients for gene/cell matrix elements.
**(B):** Mean per-cell Spearman correlation coefficient R; Mean Absolute Relative Deviation; Mean proportion of false positive genes per cell.

**(A)**

| | Simulated Truth | STARsolo | Alevin_full-decoy | Alevin_partial-decoy | Alevin_sel-align | Alevin_sketch | Kallisto |
|---|---|---|---|---|---|---|---|
| Simulated Truth | 1 | 0.951 | 0.944 | 0.879 | 0.814 | 0.614 | 0.686 |
| STARsolo | 0.951 | 1 | 0.98 | 0.902 | 0.831 | 0.623 | 0.686 |
| Alevin_full-decoy | 0.944 | 0.98 | 1 | 0.917 | 0.844 | 0.626 | 0.688 |
| Alevin_partial-decoy | 0.879 | 0.902 | 0.917 | 1 | 0.913 | 0.616 | 0.659 |
| Alevin_sel-align | 0.814 | 0.831 | 0.844 | 0.913 | 1 | 0.601 | 0.631 |
| Alevin_sketch | 0.614 | 0.623 | 0.626 | 0.616 | 0.601 | 1 | 0.645 |
| Kallisto | 0.686 | 0.686 | 0.688 | 0.659 | 0.631 | 0.645 | 1 |

**(B)**

| | mean per-cell R | MARD | false positive genes per cell |
|---|---|---|---|
| Simulated Truth | 1 | 0 | 0.0% |
| STARsolo | 0.982 | 0.047 | 1.9% |
| Alevin_full-decoy | 0.98 | 0.054 | 1.9% |
| Alevin_partial-decoy | 0.948 | 0.111 | 7.9% |
| Alevin_sel-align | 0.928 | 0.143 | 11.0% |
| Alevin_sketch | 0.831 | 0.303 | 25.1% |
| Kallisto | 0.736 | 0.349 | 30.1% |

**Figure S4:** Mapping/quantification comparison for real dataset 10X-pbmc-5k (Results 2.5).
**(A):** Matrix of Spearman correlation coefficients for gene/cell matrix elements.
**(B):** Mean per-cell Spearman correlation coefficient R; Mean Absolute Relative Deviation; Mean proportion of extra genes per cell.

**(A)**

| | CellRanger | STARsolo_sparseSA | STARsolo_fullSA | Alevin_full-decoy | Alevin_partial-decoy | Alevin_sel-align | Alevin_sketch | Kallisto |
|---|---|---|---|---|---|---|---|---|
| CellRanger | 1 | 1 | 0.998 | 0.868 | 0.808 | 0.769 | 0.759 | 0.671 |
| STARsolo_sparseSA | 1 | 1 | 0.998 | 0.868 | 0.808 | 0.769 | 0.759 | 0.671 |
| STARsolo_fullSA | 0.998 | 0.998 | 1 | 0.868 | 0.808 | 0.769 | 0.759 | 0.671 |
| Alevin_full-decoy | 0.868 | 0.868 | 0.868 | 1 | 0.93 | 0.885 | 0.752 | 0.66 |
| Alevin_partial-decoy | 0.808 | 0.808 | 0.808 | 0.93 | 1 | 0.937 | 0.714 | 0.619 |
| Alevin_sel-align | 0.769 | 0.769 | 0.769 | 0.885 | 0.937 | 1 | 0.693 | 0.598 |
| Alevin_sketch | 0.759 | 0.759 | 0.759 | 0.752 | 0.714 | 0.693 | 1 | 0.669 |
| Kallisto | 0.671 | 0.671 | 0.671 | 0.66 | 0.619 | 0.598 | 0.669 | 1 |

**(B)**

| | mean per-cell R | MARD | extra genes per cell |
|---|---|---|---|
| CellRanger | 1 | 0 | 0.0% |
| STARsolo_sparseSA | 1 | 0 | 0.0% |
| STARsolo_fullSA | 0.999 | 0.002 | 0.1% |
| Alevin_full-decoy | 0.963 | 0.119 | 1.0% |
| Alevin_partial-decoy | 0.935 | 0.165 | 6.5% |
| Alevin_sel-align | 0.92 | 0.187 | 9.1% |
| Alevin_sketch | 0.881 | 0.248 | 20.7% |
| Kallisto | 0.834 | 0.328 | 26.1% |

**Figure S5:** Real 10X-pbmc-5k dataset: clustering and neighborhood graph (Results 2.6).
**(A):** 9 cell types detected by Leiden clustering of *CellRanger*'s gene/cell matrix.
**(B):** Violin plots for cluster markers (from the *Seurat pbmc-3k* tutorial) used to identify cluster cell types.

**(A)** **(B)**

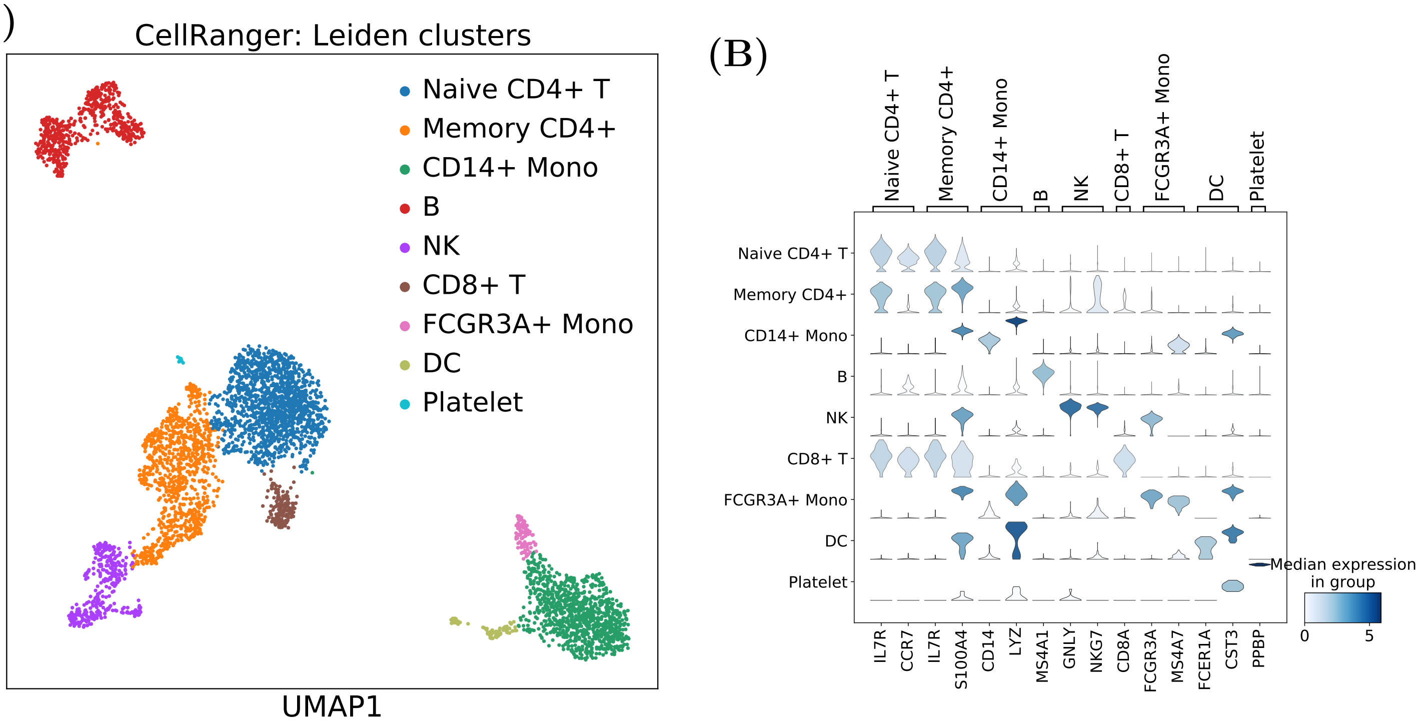

**Figure S6:** Log2-fold-changes for significantly ($p_{adj} < 0.01$) differentially expressed genes in the *Memory CD4+* cluster, each tool vs *CellRanger*, in the 10X-pbmc-5k dataset (Results 2.6). The $log_2(FC)$ values were truncated at $-10$ and 10. The genes that were not detected by a tool were assigned $log_2(FC) = 0$.

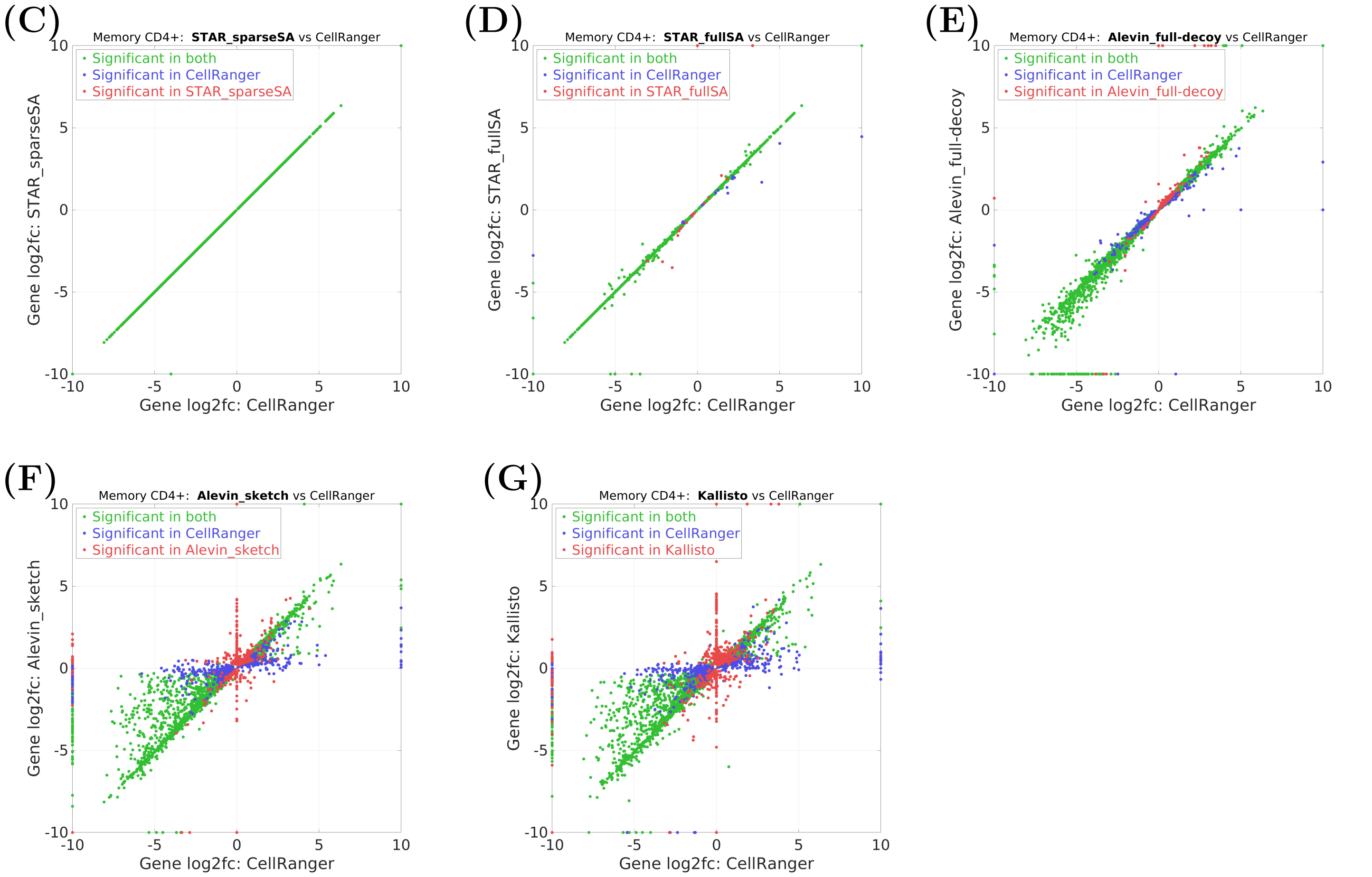

**Figure S7:** Log2-fold-changes for significantly ($p_{adj} < 0.01$) differentially expressed genes in the *CD14+ Mono* cluster, each tool vs *CellRanger*, in the 10X-pbmc-5k dataset (Results 2.6). The $log_2(FC)$ values were truncated at $-10$ and 10. The genes that were not detected by a tool were assigned $log_2(FC) = 0$.

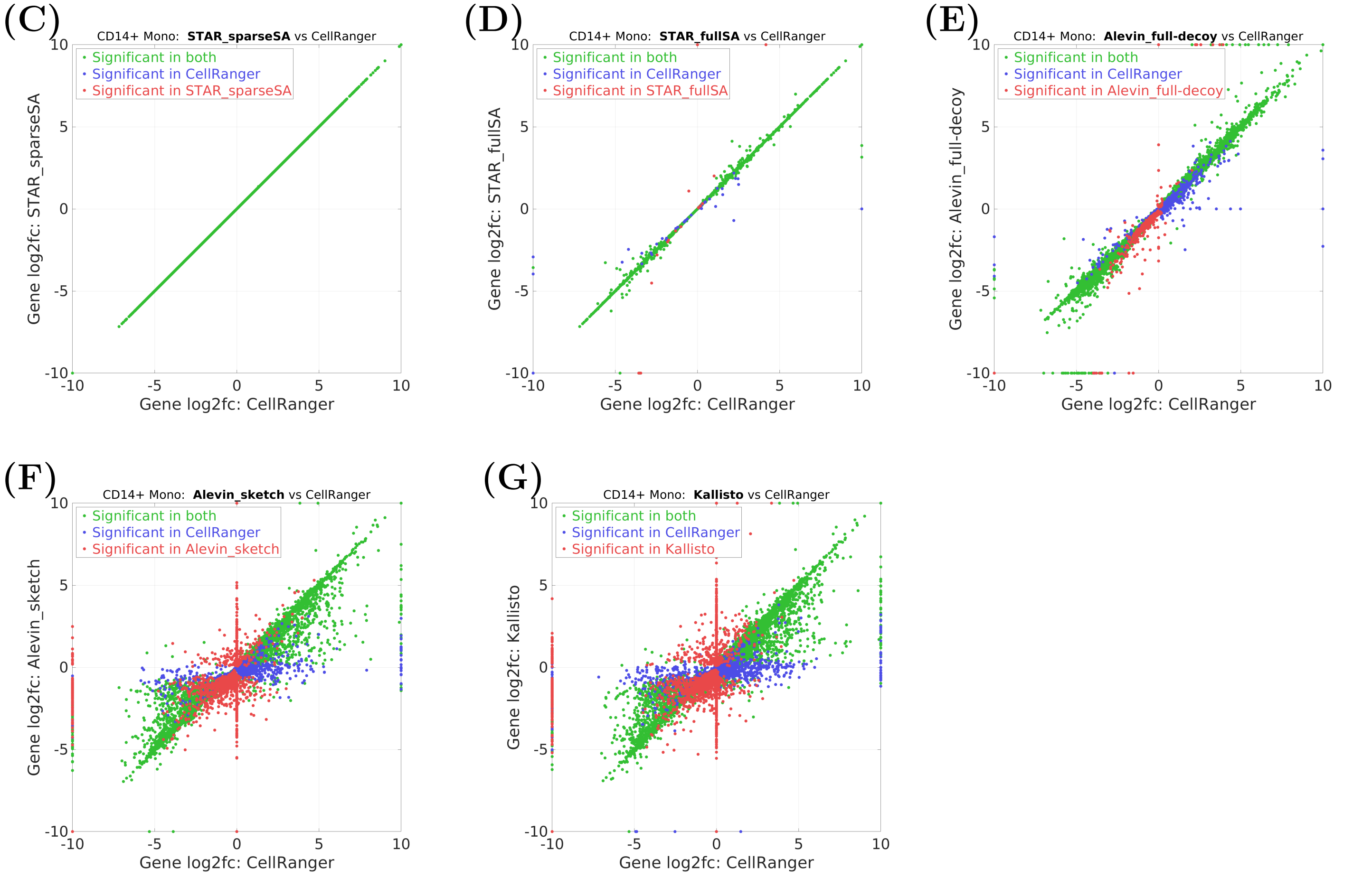

**Figure S8:** Log2-fold-changes for significantly ($p_{adj} < 0.01$) differentially expressed genes in the $B$ cluster, each tool vs *CellRanger*, in the 10X-pbmc-5k dataset (Results 2.6). The $log_2(FC)$ values were truncated at $-10$ and 10. The genes that were not detected by a tool were assigned $log_2(FC) = 0$.

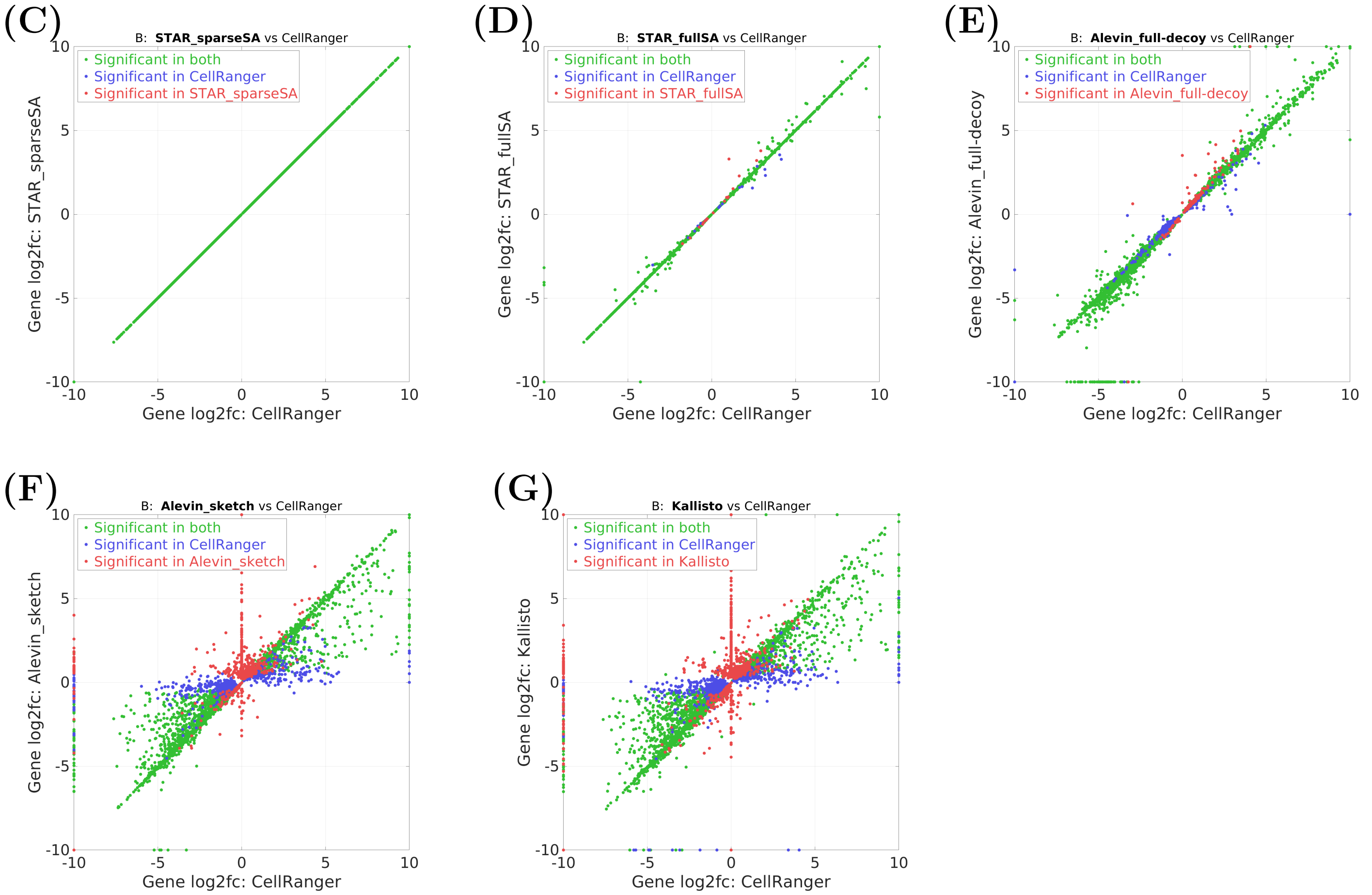

**Figure S9:** Log2-fold-changes for significantly ($p_{adj} < 0.01$) differentially expressed genes in the *NK* cluster, each tool vs *CellRanger*, in the 10X-pbmc-5k dataset (Results 2.6). The $log_2(FC)$ values were truncated at $-10$ and 10. The genes that were not detected by a tool were assigned $log_2(FC) = 0$.

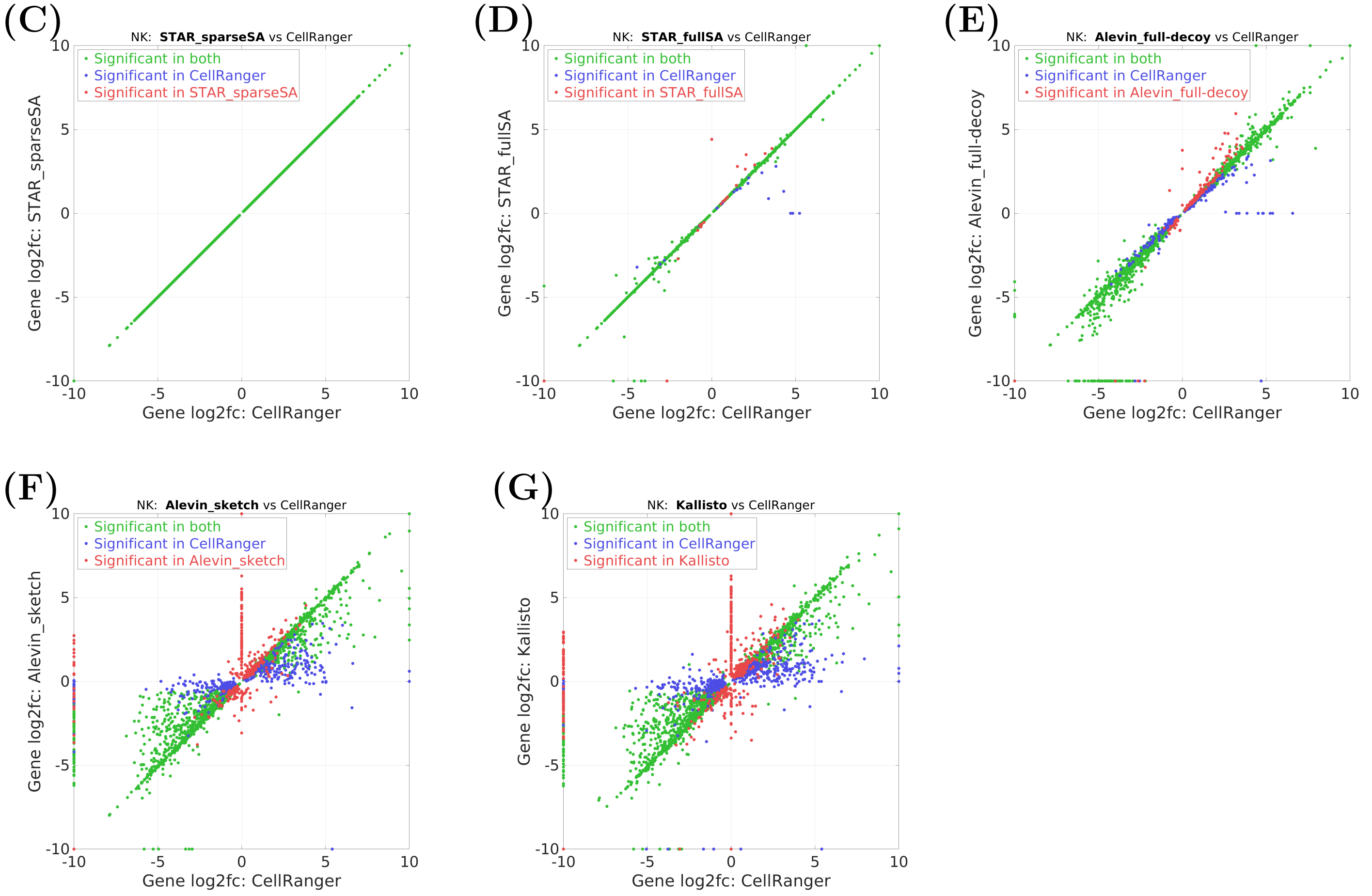

**Figure S10:** Log2-fold-changes for significantly ($p_{adj} < 0.01$) differentially expressed genes in the *CD8+ T* cluster, each tool vs *CellRanger*, in the 10X-pbmc-5k dataset (Results 2.6). The $log_2(FC)$ values were truncated at $-10$ and 10. The genes that were not detected by a tool were assigned $log_2(FC) = 0$.

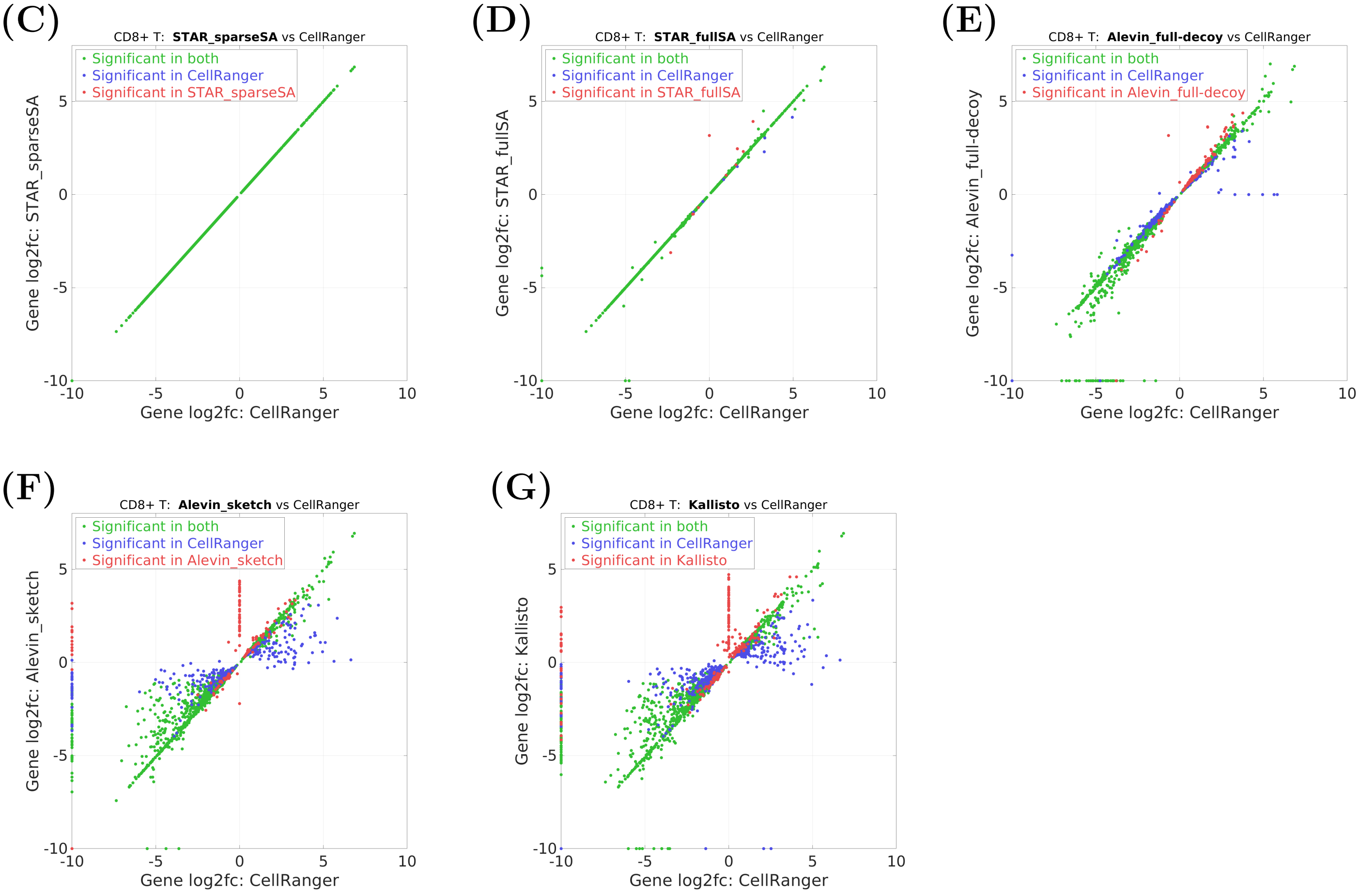

**Figure S11:** Log2-fold-changes for significantly ($p_{adj} < 0.01$) differentially expressed genes in the *FCGR3A+ Mono* cluster, each tool vs *CellRanger*, in the 10X-pbmc-5k dataset (Results 2.6). The $log_2(FC)$ values were truncated at $-10$ and 10. The genes that were not detected by a tool were assigned $log_2(FC) = 0$.

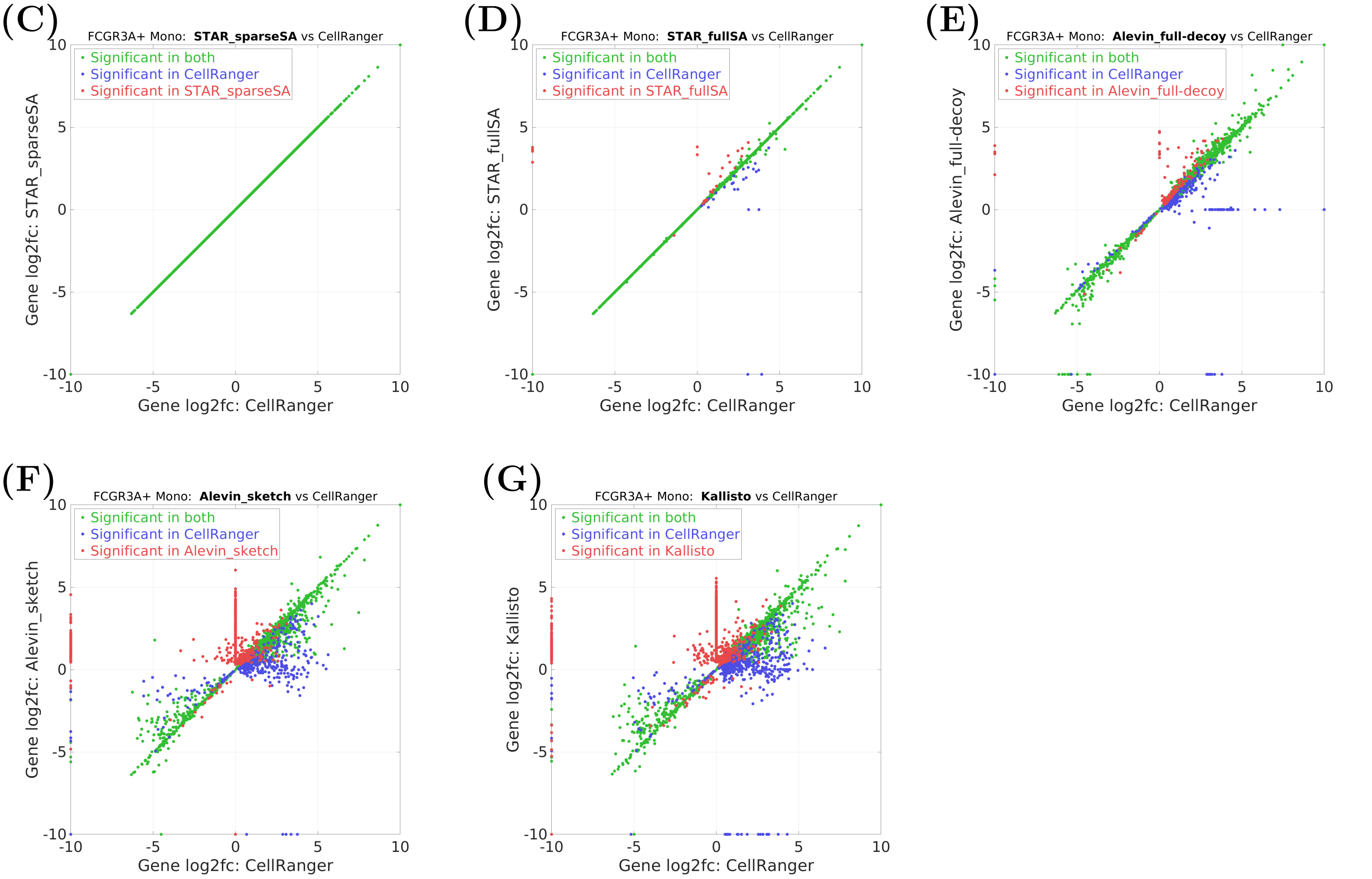

**Figure S12:** Log2-fold-changes for significantly ($p_{adj} < 0.01$) differentially expressed genes in the *DC* cluster, each tool vs *CellRanger*, in the 10X-pbmc-5k dataset (Results 2.6). The $log_2(FC)$ values were truncated at $-10$ and 10. The genes that were not detected by a tool were assigned $log_2(FC) = 0$.

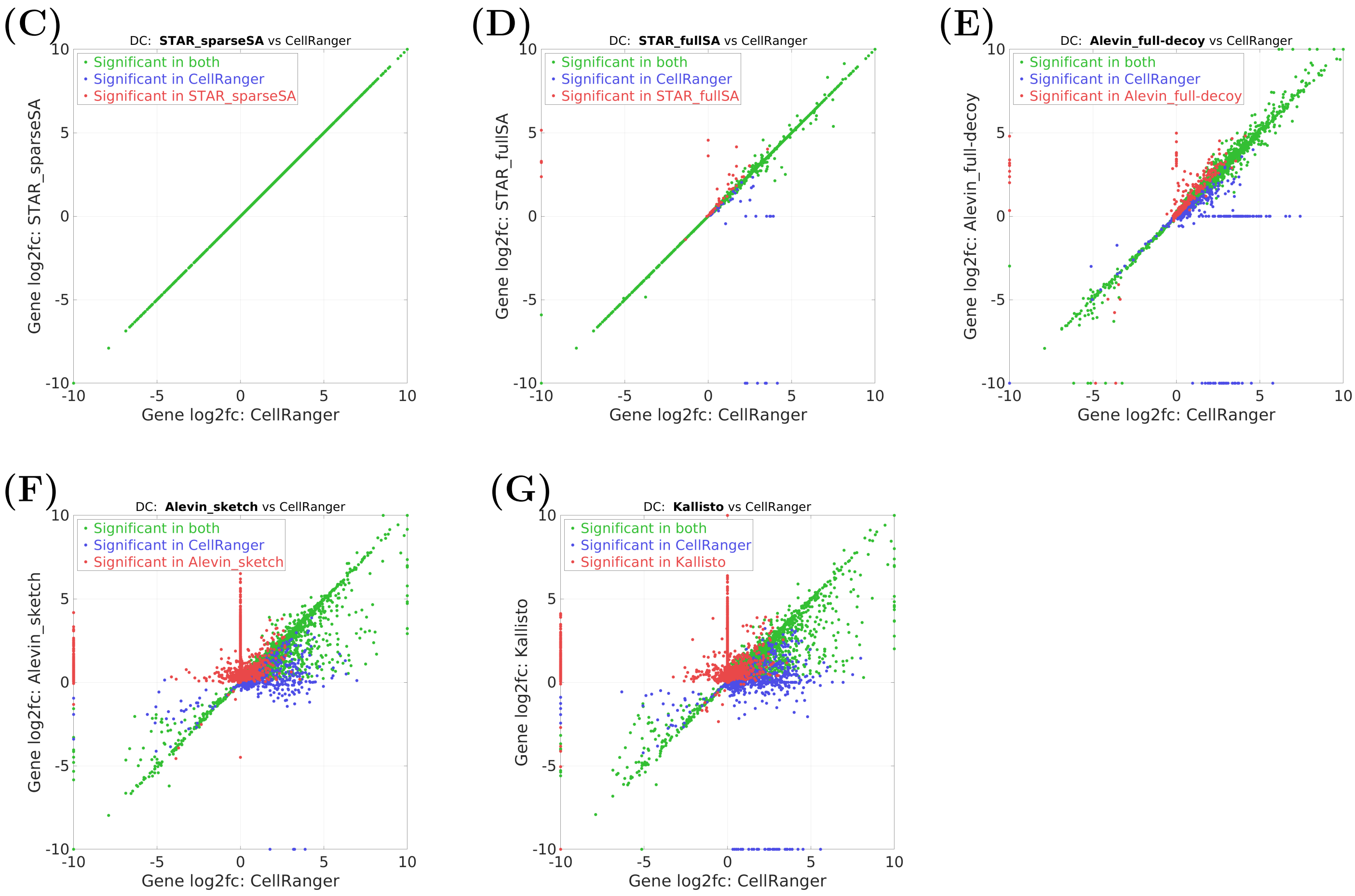

**Figure S13:** Log2-fold-changes for significantly ($p_{adj} < 0.01$) differentially expressed genes in the *Platelet* cluster, each tool vs *CellRanger*, in the 10X-pbmc-5k dataset (Results 2.6). The $log_2(FC)$ values were truncated at $-10$ and 10. The genes that were not detected by a tool were assigned $log_2(FC) = 0$.

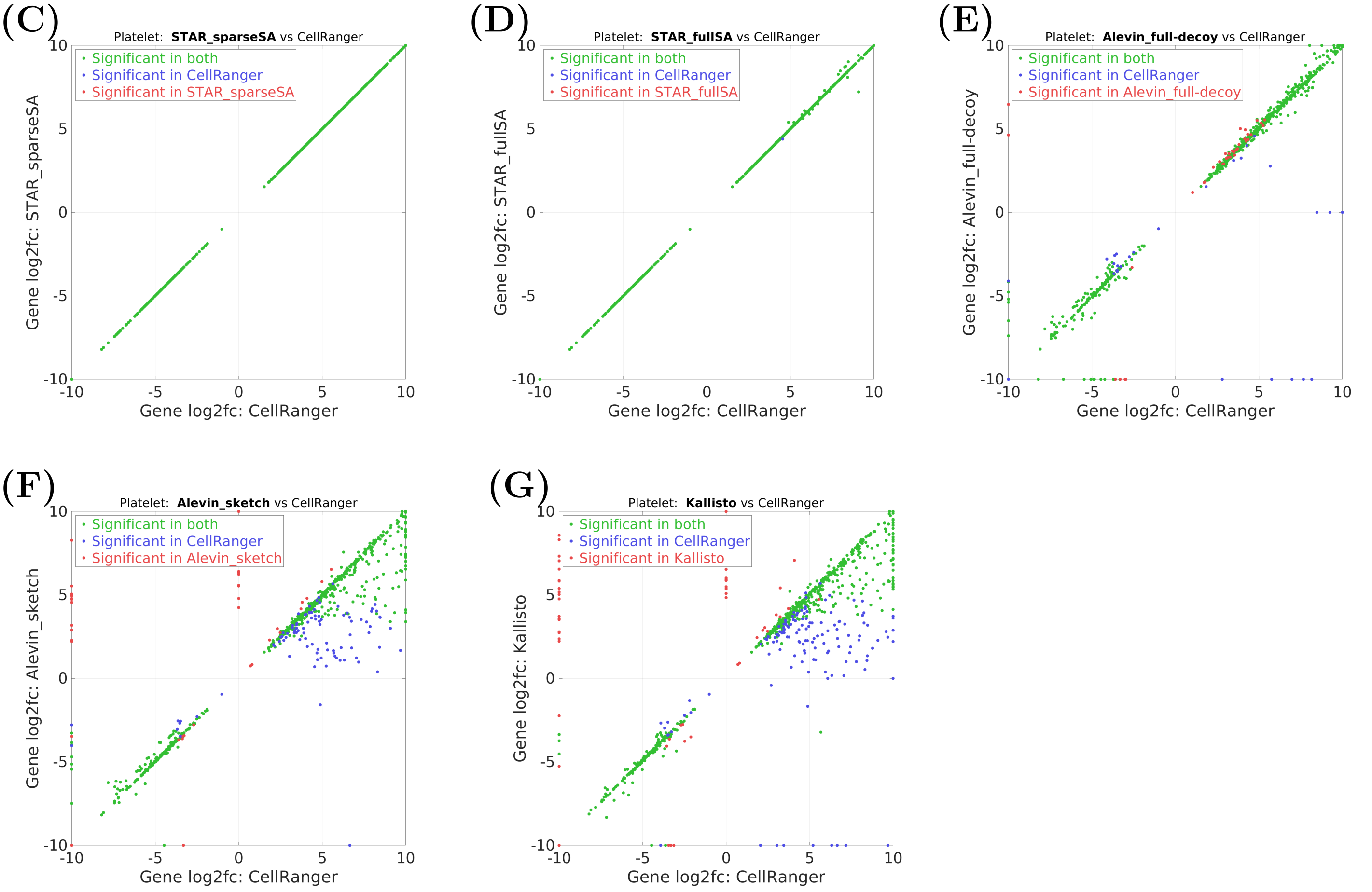

### 2 Supplementary Methods

#### 2.1 Detailed description of the splicing pipeline

The first step in this analysis was to load and parse the data from Tabula Muris and from the *STAR* genome creation. We stored the Tabula Muris annotations as a dictionary mapping from cell barcode to tissue and cell type. We also loaded the Splice Junctions that were generated by *STAR* during genome creation, along with their corresponding genes. We stored this SJ information as a dictionary mapping from SJ to a set of gene IDs. Each SJ is represented by a tuple containing the chromosome on which the SJ falls, its start and end positions, and its genomic strand. The gene IDs are numeric indices corresponding to the gene's position in the list of genes generated by *STAR* during genome creation.

After loading the fixed Tabula Muris and genome-related information, we loaded the mapping-related information. From the *STARsolo* mapping, we used the list of splice junctions found during mapping, as well as the unfiltered gene and splice junction counts. We stored both a set of all SJs seen during mapping and a list of SJs for each sequencing run, skipping SJs marked as unannotated. For SJs for which the strand was unknown, we checked the list of SJs from genome generation to see which strand(s) contained a splice junction at this position. The *STARsolo* SJ and gene counts were each concatenated for all samples along the cell barcode dimension. An additional step was required during loading of the SJ counts, which involved adding all-zero rows for SJs that were found in at least one sample but were not present in the sample being loaded. Additionally, rows corresponding to SJs that were present on both strands needed to be duplicated.

In order to facilitate our analyses, we next adjusted the splice junction and gene count matrices to align their indices. For each matrix, we added copies of rows whose corresponding SJ/gene mapped to multiple genes/SJs. Finally, we sorted both matrices according to the genomic position of the SJ indices. Ties in position, created by duplicating rows for SJs with multiple genes, were broken by the corresponding gene ID for each instance of the SJ. The end results of this adjustment step were SJ and gene count matrices with a 1:1 correspondence between their indices. We also maintained a copy of the gene count matrix before this adjustment step, to be used for later calcuations.

After preparing the data, we generated normalized splice junction expression values, denoted as U values, for each splice junction via a bootstrapping algorithm. Each iteration of the bootstrapping was performed by randomly sampling all cells with replacement, and then summing the SJ and gene counts for the selected cells and normalizing the summed SJ count by the summed gene count. For SJs with a summed gene count of zero across the selected cells, the U value was set to zero. This bootstrapping was performed for 100 iterations, and was performed separately for each cell type.

We used these bootstrapped U values to calculate log2 fold changes between cell types. Before lfc calculation, we removed all cell types that did not have more than 200 cells. For each pair of cell types within each tissue type, we calculated log2 fold change values at three different splice junction count thresholds: 50, 100, and 200. The log2 fold change values were calculated using the median of the 100 bootstrapped U values for each cell type. At this step we also applied several filters to the calculated log2 fold change values based on the un-normalized SJ and gene count matrices. The filter applied to the SJ counts was that the sum of the SJ counts for all cells in the cell type was greater than the SJ count threshold being applied. The genes were filtered to only include genes whose total cell type expression was greater than the 75th percentile of cell type gene expression for all genes. We required that the gene threshold be met in both cell types, whereas the SJ threshold was only required of at least one cell type. We removed rows from the matrix if their corresponding gene or SJ did not pass the filtering step. The remaining log2 fold change values were plotted in a histogram (Figure 2.8B), and colored according to the highest SJ threshold that they passed. We also plotted a comparison of the log2 fold change value and the maximum U value between the two cell types being compared (Figure res:splicingA).

In addition to the cell type-level plots, we also plotted aggregate plots at the tissue type level (Figures 2.8C-E)and over all tissues types (Figure 2.8F). To aggregate across cell type pairs within a given tissue type, we took the max log2 fold change value for all SJ/gene combinations. As before, we plotted a histogram of the aggregated log2 fold change values colored by the highest SJ threshold that they passed, however for the aggregated plots we plotted the absolute value of the log2 fold changes. To aggregate over all tissue types, we concatenated the log2 fold change values for all tissue types.
